# An Arf/Rab cascade controls the growth and invasiveness of glioblastoma

**DOI:** 10.1101/2020.04.30.070334

**Authors:** Gopinath Kulasekaran, Mathilde Chaineau, Valerio E. Piscopo, Federica Verginelli, Maryam Fotouhi, Martine Girard, Yeman Tang, Rola Dali, Rita Lo, Stefano Stifani, Peter S. McPherson

**Affiliations:** Department of Neurology and Neurosurgery, Montreal Neurological Institute, McGill University, Montreal, QC, Canada

## Abstract

Glioblastoma is the most common and deadly malignant brain cancer. We now demonstrate that loss of function of the endosomal GTPase Rab35 in human brain tumor initiating cells (BTICs) increases glioblastoma growth and decreases animal survival following BTIC implantation in mouse brain. Mechanistically, we identify that the GTPase Arf5 interacts with the guanine nucleotide exchange factor (GEF) for Rab35, DENND1/connecdenn and allosterically enhances its GEF activity towards Rab35. Knockdown of either Rab35 or Arf5 increases cell migration, invasiveness and self-renewal in culture and enhances the growth and invasiveness of BTIC-initiated brain tumors in mice. RNAseq of the tumors reveals upregulation of the tumor-promoting transcription factor SPOCD1, and disruption of the Arf5/Rab35 axis in glioblastoma cells leads to strong activation of the epidermal growth factor receptor with resulting enhancement of SPOCD1 levels. These discoveries reveal an unexpected cascade between an Arf and a Rab and indicate a role for the cascade, and thus endosomal trafficking, in brain tumors.

## Introduction

Vesicle trafficking is a critical regulator of cellular processes involved in cancer progression and tumorigenesis including cell proliferation and self-renewal, adhesion, migration, and invasiveness (Goldenring, 2013; Schmid, 2017). For example, following activation by ligand, tyrosine kinase receptors, such as the epidermal growth factor (EGF) receptor, are targeted to clathrin-coated pits and vesicles for internalization and transport to the endosomal system (Dikic, 2003). From there they either recycle back to the plasma membrane or are targeted to lysosomes for degradation (Dikic, 2003). These trafficking decisions determine the levels and signaling capacities of the receptors and thus control cell growth and proliferation (Vieira et al., 1996; Clague and Urbé, 2001; Wiley, 2003; Caldieri et al., 2018; Schmid, 2017). Similar trafficking pathways control other signaling receptors and cell surface proteins involved in cell adhesion and cell migration (Allaire et al., 2013; Porębska et al., 2018). Thus, alterations in membrane trafficking pathways are emerging as key to cancer progression.

Formation, transport and fusion of vesicles in the endosomal system are regulated by a plethora of proteins including Rab GTPases, which cycle between an active, GTP-bound form and an inactive GDP-bound form (Zerial and McBride, 2001). Once activated by their specific guanine nucleotide exchange factors (GEFs) (Marat et al., 2011), Rabs interact with effectors that mediate various trafficking functions. Thus, deregulation of the expression of Rab proteins can lead to altered membrane trafficking and cancer progression (Shaughnessy and Echard, 2018; Guadagno and Progida, 2019; Gopal Krishnan et al., 2020). In fact, Rabs are emerging as an important new set of drug targets in cancer (Qin et al., 2017).

Glioblastoma multiforme (GBM), or grade IV glioma, is the most aggressive tumor that begins in the brain, and due to its extreme invasive and proliferative nature, the median survival time following diagnosis is approximately 14 months (De Bonis et al., 2013; Wirsching et al., 2016). There have been several links between Rab proteins and GBM. For example, Rab27a has been linked to lysosomal exocytosis regulating glioma cell migration and invasion (Liu et al., 2012), Rab3a is highly expressed in glioma cells and human GBM samples and is involved in glioma initiation and progression (Kim et al., 2014), and the expression of Rab38 is significantly increased in GBM (Wang et al., 2013).

Rab35 has been extensively studied based on its function in the endosomal system (Chaineau et al., 2013). For example, we demonstrated that Rab35 is responsible for the recycling of cadherins from early endosomes to the plasma membrane and therefore regulates cell adhesion (Allaire et al., 2013). Furthermore, by recruiting ACAP2, a GTPase activating protein (GAP) for Arf6, activated Rab35 suppresses the recycling of β1-integrin to the cell surface, decreasing cell migration (Allaire et al., 2013; Rahajeng et al., 2012; Chesneau et al., 2012; Kobayashi and Fukuda, 2012; Miyamoto et al., 2014). Thus, loss of Rab35 function leads to decreased cell adhesion and increased cell migration, processes associated with cancer progression. Loss of Rab35 also causes enhanced recycling of EGF receptor and increased cell proliferation (Allaire et al., 2013). Consistently, we demonstrated that the mRNA level of Rab35 is decreased in resected human glioblastoma (Allaire et al., 2013). We thus sought to investigate the role of Rab35 in progression of glioblastoma.

## Results

### Disrupting Rab35 increases brain tumor growth and decreases host survival

The cellular components of GBM are highly heterogeneous (Patel et al., 2014), including cells with stem cell-like properties called brain tumor initiating cells (BTICs) (Chen et al., 2010; Couturier et al., 2020). BTICs are noted for their rapid proliferation, self-renewal, and their ability to induce the formation of tumors when transplanted into host mice (Kelly et al., 2009). We silenced the expression of Rab35 in the BTIC line BT025 (Kelly et al., 2009) using a previously described lentiviral system (Thomas et al., 2009; Ritter et al., 2017) that drives expression of GFP and shRNAmiR sequences selective for Rab35 (Allaire et al., 2013). Both sequences targeting Rab35 lead to efficient protein reduction (Fig. 1A). Control cells or cells with reduced Rab35 expression (500,000/mouse) were stereotactically implanted into the right striatum of host NOD-SCID immunodeficient mice and tumor growth was assessed through the GFP signal emanating from the human cells (Fig. 1B). At 7 days following injection, GFP-positive cells were barely detectable with the control shRNAmiR transduced BTICs whereas extensive green fluorescence was seen in the brains of host mice injected with Rab35 knockdown cells, indicating enhanced tumor growth (Fig. 1B/D). The larger tumors were also detected using H & E staining (Fig. 1C). Importantly, implantation of Rab35 knockdown cells decreases the survival of host mice compared to mice implanted with non-silenced BT025 cells (Fig. 1E). Thus, Rab35 knockdown increases tumor growth and shortens life span.

**Figure 1.**
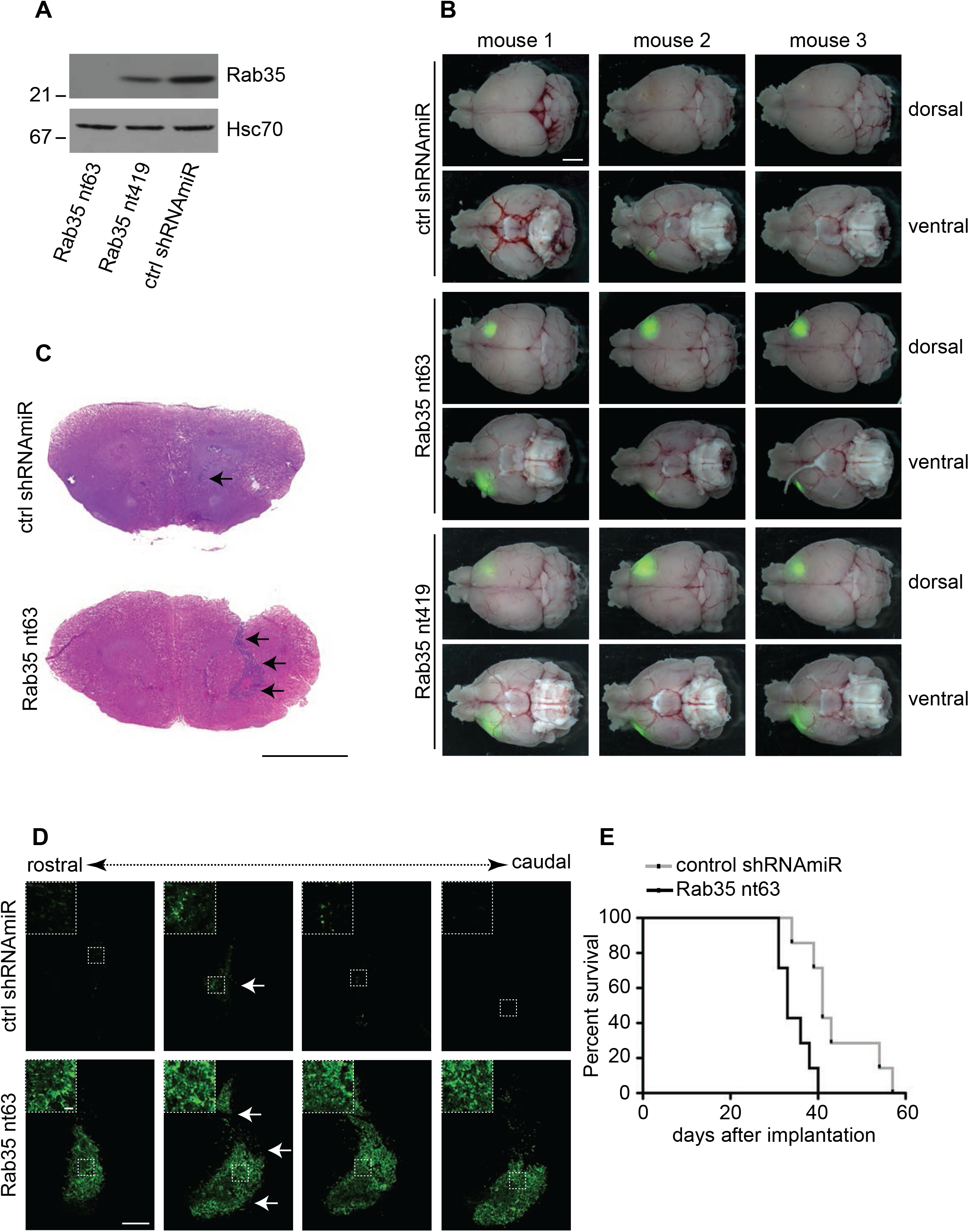
Inhibiting the expression of Rab35 increases tumor growth *in vivo*. **(A)** BT025 cells were transduced with a non-targeting control (ctrl) shRNAmiR or two different shRNAmiRs targeting Rab35 (nt63 and nt419). Cell lysates were processed for immunoblot using antibodies recognizing the indicated proteins. The migration of molecular mass markers (in kDa) is indicated. **(B)** BT025 cells (5×10^5^ cells) transduced as in **A** were stereotactically injected into the right striatum of NOD-SCID mice that were euthanized 1 week after implantation and brains were dissected. Tumor growth was monitored based on GFP fluorescence from the viral cassette. Three representative mouse brains (dorsal and ventral views) are shown for each condition. Scale bar = 2 mm. Hematoxylin and eosin (H&E) staining **(C)** and GFP staining **(D)** of coronal sections of brains of mice implanted with BT025 cells transduced with control (ctrl) or Rab35 nt63 shRNAmiRs. Scale bars = 2 mm. Arrows point to the tumors. Magnified views of the tumors in **D** are shown as insets with the scale bar = 200 μm. **(E)** Kaplan– Meier survival curves of mice implanted with BT025 cells transduced as in **A-D**. N = 7 mice for Rab35 nt63, N = 7 mice for ctrl. The statistical significance was examined by Mantel–Cox test (p=0.0029).

### Overexpression of Rab35 decreases tumor growth and prolongs survival

We next used lentiviral delivery to overexpress GFP-Rab35 in BT025 cells. Control cells were transduced with a construct expressing GFP alone. After 2 weeks, tumors observed in brains of mice implanted with the GFP-Rab35 expressing BTICs were smaller than the ones observed in brains implanted with the GFP expressing BTICs (Fig. 2A). Importantly, the survival of mice implanted with GFP-Rab35 expressing cells was prolonged in comparison with the survival of mice implanted with control GFP expressing cells for both BT025 cells (Fig. 2B) and a second BTIC line BT048 (Fig. 2C). Together, the data in figure 1 and 2 demonstrate that Rab35 expression controls GBM development *in vivo*.

**Figure 2.**
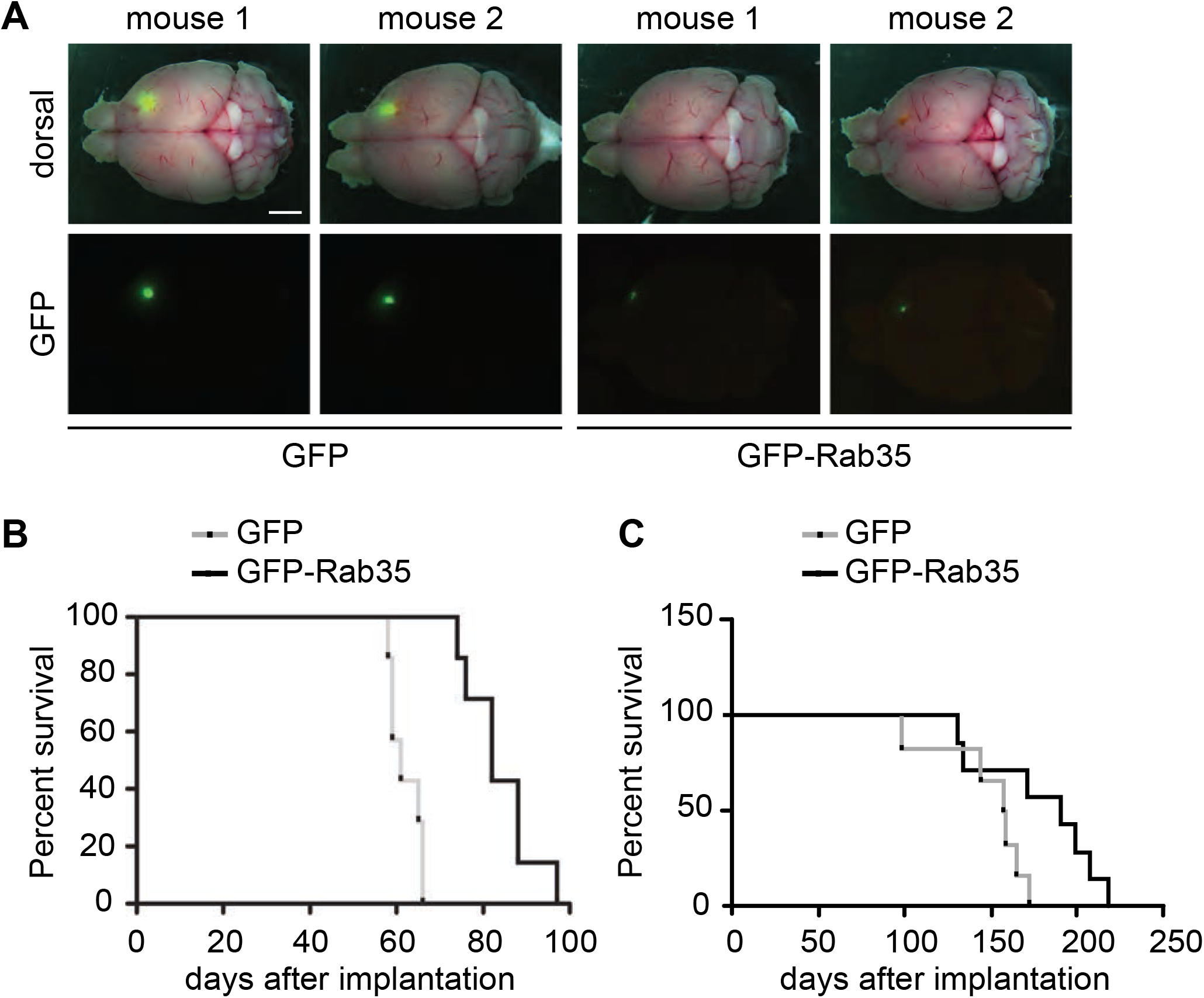
Overexpression of Rab35 limits tumor growth and lengthens lifespan. **(A)** BT025 cells (5×10^5^ cells) transduced with lentivirus expressing GFP or GFP-Rab35 were injected into the right striatum of NOD-SCID mice that were euthanized at 2 weeks after implantation. Dorsal views with (top row) and without (bottom rows) the whole-mount image are shown for two representative mice for each condition. Scale bar = 2 mm. **(B)** Kaplan–Meier survival curves of mice implanted with BT025 cells transduced as in **A** (GFP, n=7 mice; GFP-Rab35, n=7 mice). The statistical significance was examined by Mantel–Cox test (p=0.0002) **(C)** Kaplan–Meier survival curves of mice implanted with BT048 cells (5×10^5^ cells) overexpressing GFP or GFP-Rab35 (GFP, n=6 mice; GFP-Rab35, n=7 mice). The statistical significance was examined by Mantel–Cox test (p=0.055)

### The Rab35 GEF connecdenn/DENND1 binds Arf5

The levels of active Rab35 are controlled in part by its GEFs, members of the connecdenn/DENND1 family (Marat and McPherson, 2010). These proteins contain a DENN domain that harbors the GEF activity. Since: 1) Rab35 mRNA levels are reduced in GBM (Allaire et al., 2103); 2) reduction of Rab35 protein leads to enhanced growth of GBM and decreased life span (Fig. 1), and 3) Rab35 expression reduces growth and increases lifespan (Fig. 2), stimulating GEF activity towards Rab35 could provide a mechanism to improve prognosis. To identify regulators of connecdenn GEF activity, we used GST-tagged DENN domain of both connecdenn 1 and 2 in affinity-selection assays with rat brain extracts. Mass spectrometry analysis of the affinity-selected proteins revealed the presence of Arf5, a small GTPase functioning at the ER-Golgi intermediate compartment and at clathrin-coated pits during endocytosis (Chun et al., 2008; Moravec et al., 2012). The binding of Arf proteins to the DENN domains of connecdenn 1 and 2 was confirmed using a pan-Arf antibody in immunoblot on affinity-selected proteins (Fig. 3A). There are 3 classes of Arf GTPase, class II (Arf4 and 5), class I (Arf1 and 3), and class III (Arf6) (D’Souza-Schorey and Chavrier, 2006). When using Arf proteins with C-terminal GFP/CFP tags, The DENN domains of connecdenn 1 and 2 appear to bind specifically to class II Arfs with no little or no interaction with class I or III Arfs (Fig. 3B-D). Binding is strongest to Arf5, a protein that like connecdenn 1 and 2 is localized in part to clathrin-coated pits (Moravec et al., 2012). This is meaningful because connecdenn 1 and 2 and Rab35 also localize to these endocytic structures (Allaire et al., 2010). Binding to purified Arf5 without a tag is also detected (Fig. S1A-C). Thus, while we cannot rule out that other Arf GTPase have meaningful interactions with the DENN domains, they appear to interact most prominently with class II Arfs including Arf5.

**Figure 3.**
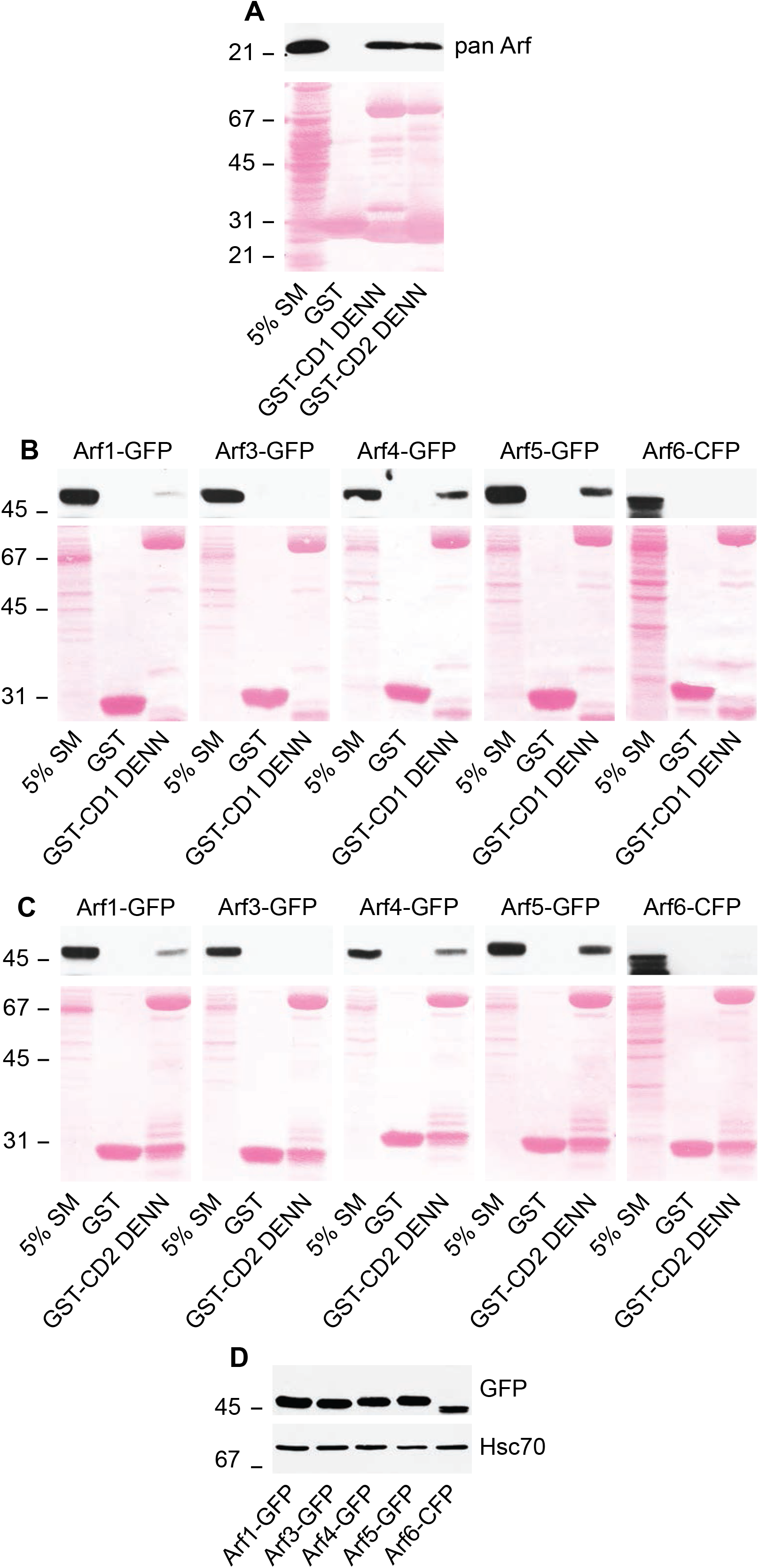
Arf5 binds the connecdenn DENN domain. **(A)** A detergent soluble lysate was prepared from adult rat brain and 2 mg of total protein was incubated with GST or GST conjugated to the DENN domain of connecdenn 1 (CD1) and 2 (CD2), pre-bound to glutathione-Sepharose. The glutathione-Sepharose beads were washed and bound proteins were prepared for immunoblot with a pan Arf antibody. An aliquot of the brain lysate equal to 5% of that added to the beads was run as a starting material (SM). The ponceau stained transfer indicates the levels of fusion protein. The migration of molecular mass markers (in kDa) is indicated. **(B/C)** Detergent soluble lysates were prepared from HEK-293 cells transfected with various Arf isoforms with C-terminal GFP or CFP tags as indicated. Soluble lysates were prepared form the cells and 0.4 mg were incubated with GST alone or GST conjugated to the DENN domain of connecdenn 1 (CD1) **(B)** and 2 (CD2) **(C)**. The glutathione-Sepharose beads were washed and the bound proteins were prepared for immunoblot with an antibody recognizing GFP. An aliquot of the cell lysate equal to 5% of that added to the beads was run as a starting material (SM). The ponceau stained transfer indicates the levels of fusion protein. The migration of molecular mass markers (in kDa) is indicated. **(D)** Lysates from cells transfected with the indicated Arf-GFP/CFP constructs were processed for immunoblot with an antibody recognizing GFP. The migration of molecular mass markers (in kDa) is indicated.

### Arf5 interaction allosterically activates connecdenn GEF activity towards Rab35

DENN domains function as GEFs for Rab proteins (Marat et al., 2011) but may have GEF activity towards other small GTPases. As the DENN domain of connecdenn binds Arf5, we tested for GEF activity of connecdenn towards this small GTPase. Consistent with the possibility of GEF activity towards Arf5, the DENN domain of connecdenn binds Arf5 directly (Fig. S1A-C). We thus sought to explore if Arf5 binding occurs in the enzymatic pocket as seen for Rab35. A crystal structure of connecdenn 2 DENN domain allowed for the identification of 7 key residues required for Rab35 binding and GEF activity (Wu et al., 2011) (Fig. 4A). We thus generated two mutant forms of the DENN domain in which we mutated 4 key residues required for Rab35 switch I binding or 3 key residues required for switch II binding (Fig. 4A) (Wu et al., 2011). Consistent with previous reports, both sets of mutations abolish Rab35 binding, but interesting, neither influence Arf5 binding (Fig. 4B), indicating that Arf5 does not bind to the catalytic site, inconsistent with a potential GEF activity towards Arf5. Identical results were seen with mutations in the DENN domain of connecdenn 1, in which 6 of the 7 key residues are identical with one highly conserved (Fig. S1D/E). Moreover, GEFs bind their substrate GTPases in the GDP-bound form but we observed no selectivity for the DENN domains of connecdenn 1 or 2 toward the GDP-bound form of Arf5 (Fig. S1F-I). Thus, it seems unlikely that the DENN domain of connecdenn 2 has GEF activity towards Arf5.

**Figure 4.**
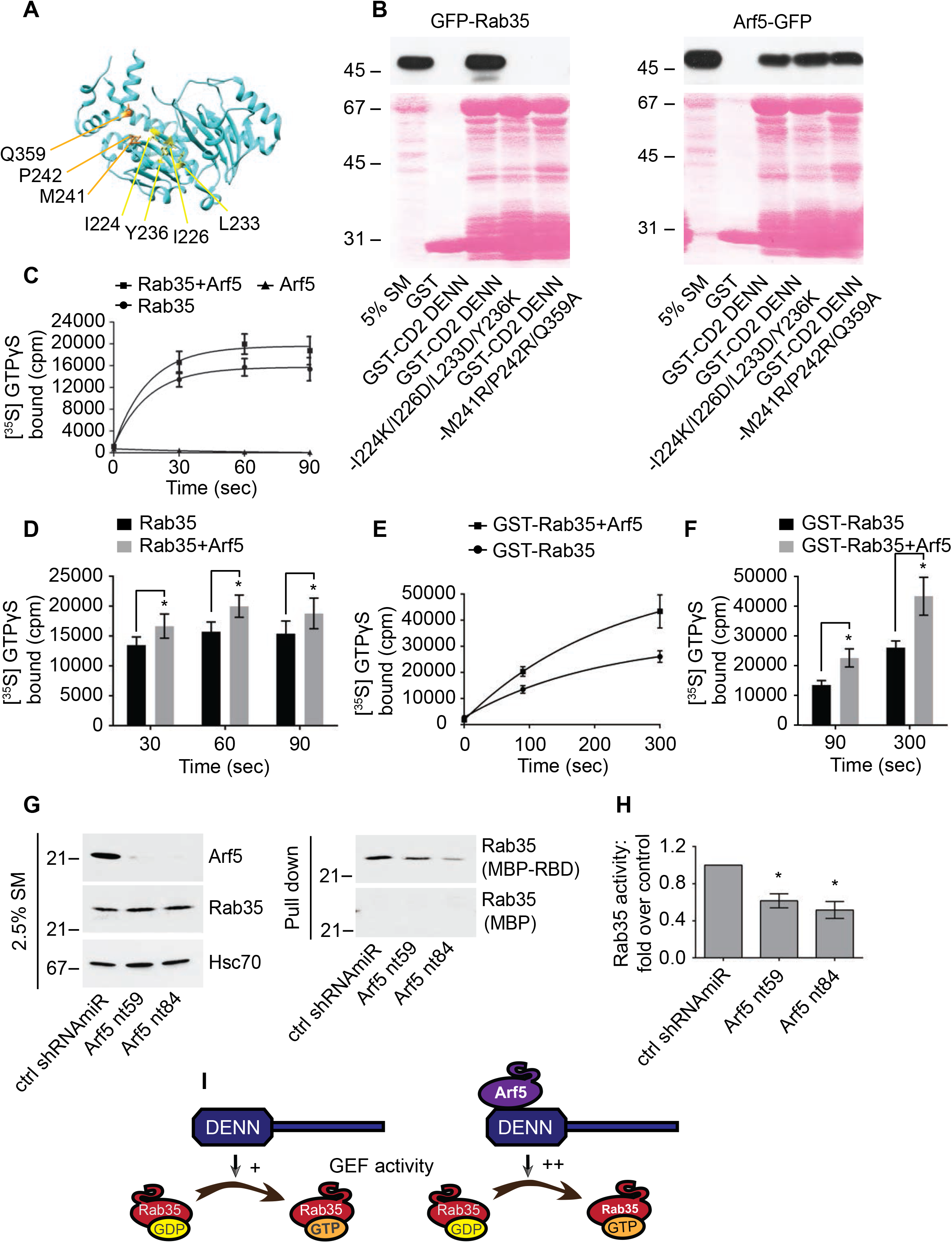
Arf5 activates connecdenn GEF activity towards Rab35. **(A)** A ribbon representation of the structure of the DENN domain of conncedenn 2 (DENND1B) taken from (Wu et al., 2011). Amino acids required for interaction with and GEF activity towards Rab35 are indicated. **(B)** Detergent soluble lysates were prepared from HEK-293 cells transfected with GFP-Rab35 or Arf5-GFP as indicated and incubated with GST alone, GST-connecdenn 2 (CD2) DENN domain, or the CD2 DENN domain with the quadrupole or triple point mutations indicated. Glutathione-Sepharose beads were washed and the bound proteins were prepared for immunoblot with an antibody recognizing GFP. An aliquot of the cell lysate equal to 5% of that added to the beads was run as a starting material (SM). The ponceau stained transfer indicates the levels of fusion protein. The migration of molecular mass markers (in kDa) is indicated. **(C)** *In vitro* GEF assays using purified connecdenn 1 (CD1) DENN domain and different combinations of Rab35 and Arf5 as indicated. The amount of [^35^S]GTPγS transferred to the GTPases was determined by collecting the reactions on filters followed by scintillation counting. The relative incorporation of [^35^S]GTPγS is plotted over time and data represent mean ± s.e.m, N=8. The curve was fit by nonlinear regression one-phase association. **(D)** Quantification of the amount of [^35^S]GTPγS transferred to Rab35 at 30, 60, and 90 sec from experiments as in **C**. Statistical analysis employed a two-way ANOVA followed by a Bonferroni’s multiple comparisons test. * p<0.05, N=8. **(E)** I*n vitro* GEF assays as described in **C** except using GST-tagged Rab35 coupled to glutathione-Sepharose beads, in the presence or absence of purified Arf5, as indicated. The amount of [^35^S]GTPγS transferred to Rab35 was determined by pelleting the beads in a microfuge. The relative incorporation of [^35^S]GTPγS is plotted over time and data represent mean ± s.e.m, N=4. The curve was fit by nonlinear regression one-phase association. **(F)** Quantification of the amount of [^35^S]GTPγS transferred to Rab35 at 90and 300 sec from experiments as in **E**. Data are shown as mean ± s.e.m. Statistical analysis employed a two-way ANOVA followed by a Bonferroni’s multiple comparisons test. * p<0.05, N=4. **(G)** Soluble lysates were prepared from HEK-293T cells transduced with control (ctrl) shRNAmiR or two different shRNAmiRs targeting Arf5 as indicated. The lysates were incubated with either MBP alone or MBP conjugated to the Rab35-binding domain of RUSC2 (MBP-RBD), with the MBP proteins pre-bound to amylose-Sepharose beads. Beads were washed and the bound proteins were processed for immunoblot with antibody specific for Rab35 (Pull down). An aliquot of the cell lysate (starting material, SM) equal to 2.5% of that added to the beads was run in parallel and immunblotted with antibodies specific for the indicated proteins. The migration of molecular mass markers (in kDa) is indicated **(H)** Quantification of the amount of Rab35 bound to the MBP-RBD from the Arf5 knockdown cells normalized to that from the control shRNAmiR-treated cells as in **G**. Data are shown as mean ± s.e.m. Statistical analysis employed a one-way ANOVA followed by a Dunnett’s post hoc test. * p<0.05, N=5. **(I)** Schematic model showing Arf5 interacts with the DENN domain of connecdenn 1/2 to stimulate its GEF activity towards Rab35.

We performed GEF assays with purified connecdenn 1 DENN domain using purified Rab35 and Arf5 as substrates. We detected no GEF activity towards Arf5, whereas activity towards Rab35 was robust (Fig. 4C). We next sought to examine if Arf5 binding could influence GEF activity towards Rab35. The Ras-specific GEF son of sevenless possess two Ras-binding sites, one at the catalytic site and a second distinct site that when bound to Ras allosterically activates Ras exchange through the catalytic site (Margarit et al., 2003). Interestingly, when we combined Arf5 with Rab35 we observed an enhancement in the GEF activity of the connecdenn DENN domain towards Rab35 (Fig. 4C/D). To confirm that the enhanced activity was towards Rab35 and not an unmasked cryptic activity towards Arf5, we repeated the GEF assays using GST-tagged Rab35 with a GST pull-down to selectively count [^35^S]GTPγS incorporated into Rab35 (the previous assay collected the GTPases on a filter). This assay confirmed that Arf5 enhanced GEF activity towards Rab35 (Fig. 4E-F). Moreover, there is no interaction between Arf5 and Rab35 (Fig. S2A/B) and Arf5 does not appear in the Rab35 pull-down from the enzyme reaction mixture (Fig. S2C/D). Thus, Arf5 binds the DENN domain of connecdenn 1 and enhances its GEF activity towards Rab35.

To test if Arf5 activates Rab35 in cells, we performed an effector-binding assay using a maltose-binding protein (MBP) fusion with the Rab35-binding domain (RBD) of RUN and SH3 domain-containing 2 (MBP-RBD), which binds Rab35 selectively in the GTP-bound form (Kobayashi et al., 2015). MBP-RBD was incubated with lysates from cells transduced with lentivirus driving Arf5 targeted shRNAmiRs. These constructs lead to loss of Arf5 without influencing the level of Rab35 (Fig. 4G). In the Arf5 knockdown cells there is a significant decrease in the binding of active Rab35 to MBP-RBD when compared to cells transduced with a control lentivirus (Fig. 4G/H). Taken together, these data demonstrate that Arf5 interaction with the connecdenn DENN domain activates its GEF activity towards Rab35 (Fig. 4I).

### The Arf5/Rab35 axis controls cell migration, invasion, and self-renewal

Enhanced cell migration and invasion are hallmarks of cancer cells. We used lentiviral-delivered shRNAmiRs to knockdown Rab35 and Arf5 and tested for alterations in these phenotypes. Treatment with virus driving shRNAmiRs directed towards Rab35 or Arf5 reduced the levels of the respective GTPases in COS-7 cells (Fig. 5A). Knockdown of either protein enhanced the migration of cells into a scratch made in a confluent monolayer of cells (Fig. 5B/C). Arf5 activates the DENN domain of connecdenn to allosterically activate its GEF activity towards Rab35 (Fig. 4C-F), placing Rab35 downstream of Arf5 in the axis (Fig. 4I). As such, expression of Rab35 may be able to rescue defects resulting from Arf5 knockdown. In fact, expression of Rab35 (Fig. S3A) rescues defects in cell migration resulting from Arf5 knockdown (Fig. 5D/E). To ensure that the observed data represented alterations in cell migration, we performed single cell tracking analysis as previously described (Ioannou et al., 2015). Both Rab35 and Arf5 knockdown increased both the distance migrated and the velocity of migration in U87 glioma cell lines (Fig. S3B/C).

**Figure 5.**
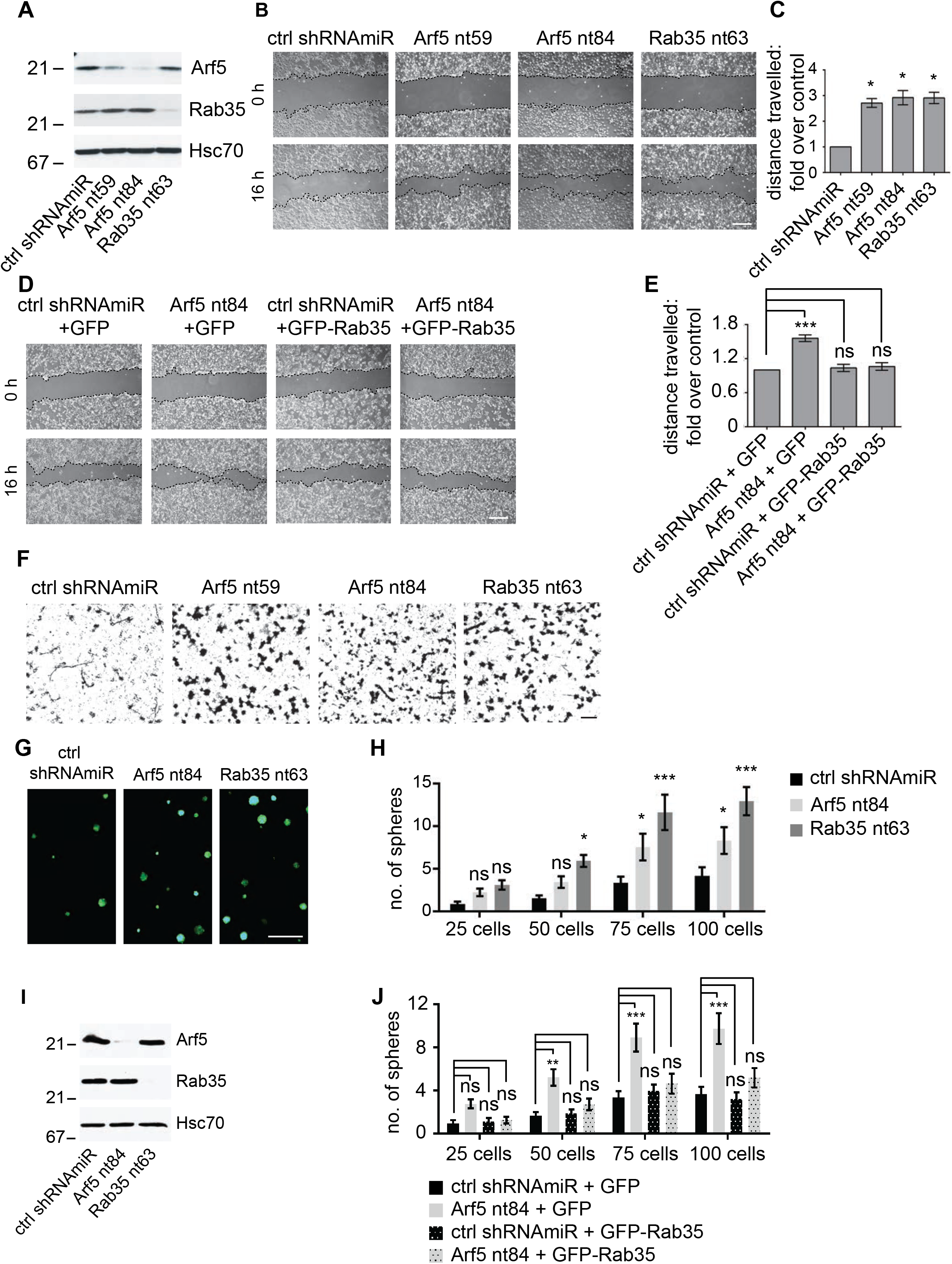
Disruption of the Arf5/Rab35 axis enhances cell migration, invasion, and self-renewal. **(A)** COS-7 cells were transduced with a control (ctrl) shRNAmiR, two different shRNAmiRs targeting Arf5, or an shRNAmiR targeting Rab35, as indicated. Crude lysates were prepared and processed for immunoblot using antibodies recognizing the indicated proteins. The migration of molecular mass markers (in kDa) is indicated. **(B)** COS-7 cells were transduced as in **A** and when confluent, a scratch was made using a pipette tip and the images were obtained immediately after (0 h) and 16 h following the scratch. Scale bar = 200 μm. **(C)** Quantification of the distance of the cells travelled into the scratch in the knockdown conditions was quantified and normalized to that seen in the control shRNAmiR from experiments as in **B**. Data are shown as mean ± s.e.m. Statistical analysis employed a one-way ANOVA followed by a Dunnett’s post hoc test. * p<0.05, N=3. **(D)** COS-7 cells were transduced with a control (ctrl) shRNAmiR plus GFP, an shRNAmiR targeting Arf5 (nt84) plus GFP, ctrl shRNAmiR plus GFP-Rab35, and an shRNAmiR targeting Arf5 (nt84) plus GFP-Rab35, as indicated. When confluent, a scratch was made using a pipette tip and the images were obtained immediately after (0 h) and 16 h following the scratch. Scale bar = 200 μm. **(E)** Quantification of the distance the cells travelled into the scratch was quantified and normalized to that seen in the control shRNAmiR plus GFP from experiments as in **D**. Data are shown as mean ± s.e.m. Statistical analysis employed a one-way ANOVA followed by a Bonferroni’s multiple comparisons test. * p<0.05, N=4. **(F)** COS-7 cells transduced as in **A** were platted on Matrigel coated transwell permeable supports with 8 μm pores in a transwell chamber. After 14 h, cells on the filter in the upper well were scrapped off the filter and the cells facing the lower chamber were fixed and stained with crystal violet. The scale bar = 50 μm. **(G)** BT025 cells were transduced with a control shRNAmiR (ctrl) or shRNAmiRs targeting Arf5 or Rab35. Cells were than counted and 100 cells were plated per well in 96-well plates and allowed to grow for 10 days. Resulting neurospheres were imaged using the GFP signal driven from the viral cassette. The scale bar = 500 μm. **(H)** Virally transduced cells as in **G** were diluted to 25, 50, 75 or 100 cells/well and the number of spheres was determined after 10 days. Data are shown as mean ± s.e.m. Statistical analysis employed a two-way ANOVA followed by a Bonferroni’s multiple comparisons test, N=3, * p<0.05, *** p<0.001, ns = not significant. **(I)** Lysates prepared from cells transduced as in **G** were processed for immunoblot with antibodies recognizing the indicated proteins. The migration of molecular mass markers (in kDa) is indicated. **(J)** BT048 cells were transduced with a control (ctrl) shRNAmiR or shRNAmiR targeting Arf5. After a week, the spheres were dissociated and the cells were transduced with either GFP or GFP-Rab35 expressing lentivirus. The transduced cells were diluted to 25, 50, 75 or 100 cells/well and the number of spheres was determined after 12 days. Data are shown as mean ± s.e.m. Statistical analysis employed a two-way ANOVA followed by a Bonferroni’s multiple comparisons test, N=4, ** p<0.01, *** p<0.001. ns = not significant.

We next examined for alterations in invasion, in which cells were plated in the upper well of a transwell chamber on filters with 8 μm pores coated with Matrigel (Ioannou et al., 2015). To migrate towards serum in the bottom chamber, cells must degrade the Matrigel and transform morphologically in order to squeeze through the pores. Both Arf5 and Rab35 knockdown enhance cell invasion (Fig. 5F). Thus, the Arf5/Rab35 axis enhances both the migratory and invasive capacities of cells.

BTICs have the inherent property to self-renew and grow into neurospheres and self-renewal is a hallmark of tumor development (Kelly et al., 2009). To study their self-renewal abilities, BTICs were plated at 25-100 cells per well and were monitored for neurosphere formation. In both BT025 (Fig. 5G-I) and BT048 (Fig. S3D-F) cells, knockdown of both Arf5 and Rab35 leads to enhanced self-renewal as observed by increased numbers of neurospheres when compared to control cells. Moreover, the enhanced self-renewal resulting from Arf5 knockdown is rescued by expression of Rab35 (Fig. 5J; Fig. S3G/H), confirming that Rab35 is functionally downstream of Arf5 in an axis that controls self-renewal of BTICs.

### Arf5 and Rab35 control the growth and migration of brain tumors

Knockdown of Rab35 in BT025 BTICs enhances tumor growth when implanted in the brains of NOD-SCID mice (Fig. 1). Given that Arf5 cascades to Rab35 we wondered if similar results would be seen with Arf5 knockdown. We thus attenuated the expression of both GTPases in BT025 cells (Fig. 6A) and stereotactically implanted the cells into the striatum. Unlike in figure 1/2 where we implanted 500,000 cells per animal, here we implanted 30,000 cells per animal. The experiments in figure1 and 2 were designed to test for the possibility of altered tumor growth and to examine survival. Large numbers of cells were used to shorten the time spans needed to generate the survival curves. The experiments in Figs. 6 and 7 were designed to more quantitatively measure tumor growth and to examine for potential alterations in cell migration. Thus, smaller numbers of cells were used to allow for slower development of the tumor, providing more controlled growth. At 4 weeks after implantation, tumors were approximately 2-times larger following Arf5 knockdown when compared to a control shRNAmiR (Fig. 6B/C) and as much as 8-times larger following Rab35 knockdown (Fig. 6B/D). Thus, loss of either Arf5 or Rab35 enhances tumor growth.

**Figure 6.**
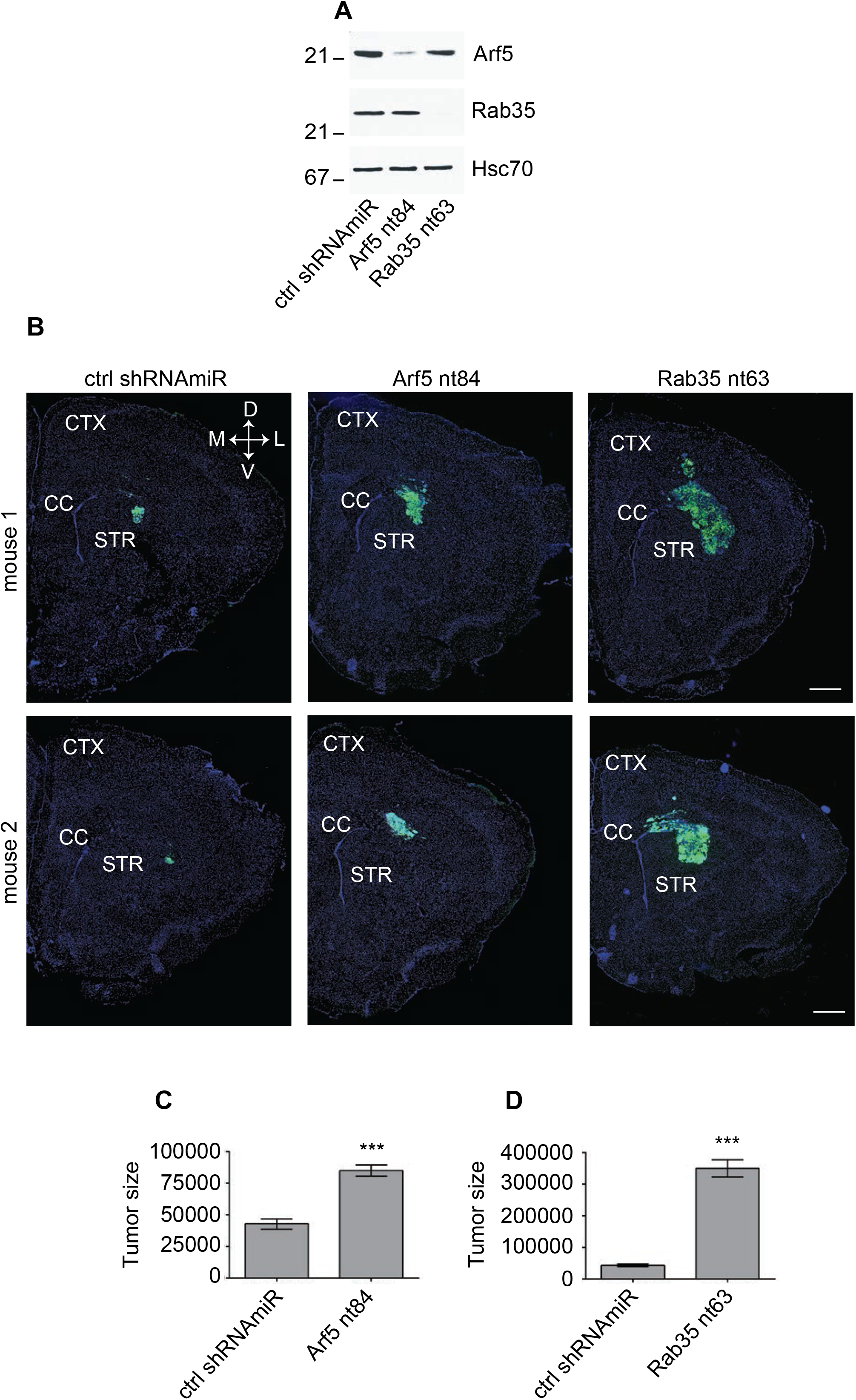
Knockdown of Arf5 or Rab35 increases tumor growth. **(A)** BT025 cells were transduced with control (ctrl) shRNAmiR or shRNAmiRs targeting Arf5 or Rab35 as indicated. Lysates prepared from cells were processed for immunoblot with antibodies recognizing the indicated proteins. The migration of molecular mass markers (in kDa) is indicated. **(B)** BT025 cells (3×10^4^ cells) transduced as in **A** were stereotactically injected into the right striatum of NOD-SCID mice, and the mice were sacrificed after 4 weeks. Dissected brains were cryosectioned in the coronal plane and the human cells were revealed through the GFP signal. Representative sections from 2 mice for each condition are shown. CTX=cortex, STR=striatum, CC=corpus callosum. Dorsal (D), ventral (V), lateral (L) and medial (M). Scale bars = 500 μm. **(C)** Serial coronal sections from mice prepared as in **B** were imaged to include all fields containing the entire injected hemisphere. The area of the tumor on each section was determined and the signal from all section was averaged to determine a tumor size in pixels. Data are shown as mean ± s.e.m. in which n=4 for control mice and n=5 for mice with Arf5 or Rab35 KD. Statistical analysis employed an unpaired t-test, ***p<0.001. **(D)** Quantification performed as in **C** but from mice injected with cells transduced with control (ctrl) shRNAmiR or shRNAmiR targeting Rab35. Note that the control in **C** and **D** are the same data. Statistical analysis employed an unpaired t-test, *** p<0.001.

**Figure 7.**
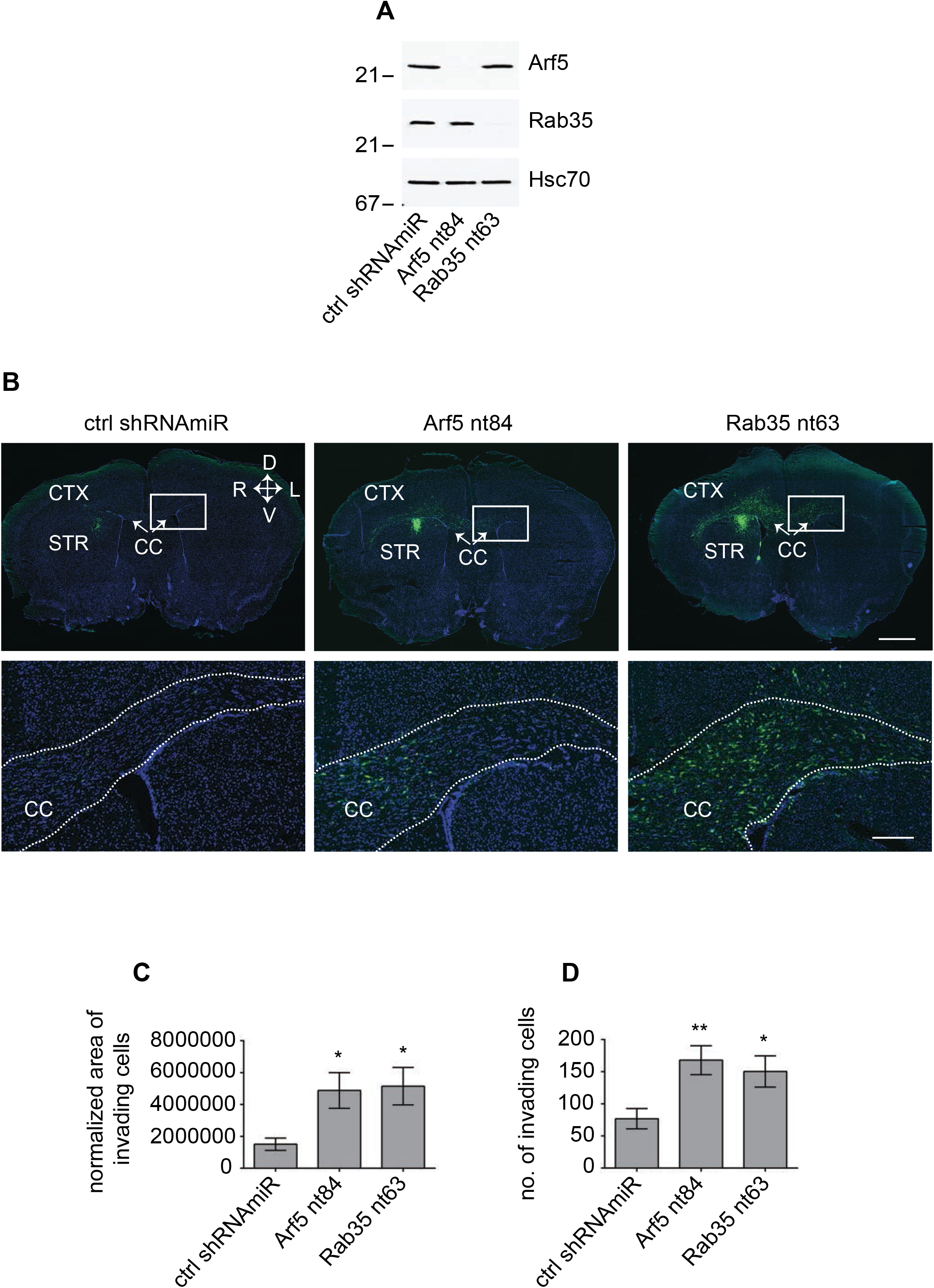
Knockdown of Arf5 or Rab35 increases tumor invasion. **(A)** BT048 cells were transduced with control (ctrl) shRNAmiR or shRNAmiRs targeting Arf5 or Rab35 as indicated. Lysates were processed for immunoblot with antibodies recognizing the indicated proteins. The migration of molecular mass markers (in kDa) is indicated. **(B)** BT048 cells (1×10^5^ cells), transduced as in **A** were stereotactically injected into the right striatum of NOD-SCID mice. Mice were euthanized 5 weeks after implantation of the cells. Brains were cryosectioned in the coronal plane and human cells were observed through the GFP signal. A representative section for each viral condition is shown. CTX=cortex, STR=striatum, CC=corpus callosum. Dorsal (D), ventral (V), left hemisphere (L) and right hemisphere (R) are indicated. The lower panels are magnified views of the boxed areas in the upper panels and come from the contralateral side from the injection. Scale bar = 1000 μm for the lower magnification and 200 μm for the higher magnification. **(C)** Serial coronal sections from mice prepared as in **B** were imaged to include all fields with human cells. Invasion of the implanted cells was calculated by determining the ratio of area of the tumor on each section over the area of invaded cells on the contralateral side to the injection. Data are shown as mean ± s.e.m. in which n=6 for control mice and n=7 for mice with Arf5 or Rab35 knockdown. Statistical analysis employed one-way ANOVA followed by a Dunnett’s multiple comparisons test, * p<0.05. **(D)** Serial coronal sections from mice prepared as in **B** were imaged to include all fields with GFP-positive cells. The number of cells on the contralateral side to the injection was determined. Data are shown as mean ± s.e.m. in which n=6 for control mice and n=7 for mice with Arf5 or Rab35 knockdown. Statistical analysis employed one-way ANOVA followed by a Dunnett’s multiple comparisons test, ** p<0.01, * p< 0.05.

Following surgical removal GBMs generally recur at a second site, including on the contralateral hemisphere of the brain from the initial tumor site (Stupp et al., 2005; Sherriff et al., 2013). Glioma cells invade the brain preferentially along white matter tracts (Pedersen et al., 1995; Beliën et al., 1999). We sought to examine if disruption of the Arf5/Rab35 axis also influenced the migration of tumor cells as part of an invasion process. Whereas BT025 cells grow as a single tumor mass (Fig. 6B), BT048 cells have enhanced migratory abilities (Verginelli et al., 2013) and grow as more disperse tumors (Fig. S4). We thus performed knockdown of Arf5 and Rab35 in BT048 cells (Fig. 7A) and stereotactically implanted 100,000 cells into the right striatum (Fig. 7B). After 5 weeks we examined for the presence of the implanted cells on the contralateral side. Disruption of both GTPases enhanced cell migration to the contralateral side (Fig. 7B), measured either by determining the area covered by the invading cells (Fig. 7C) or the number of invading cells on the contralateral side (Fig. 7D). Thus, disruption of the Arf5/Rab35 axis leads to enhanced growth and migration of brain tumors.

### SPOCD1 downstream of Rab35 and activated EGF receptor contributes to the growth of glioblastoma

We next sought to identify a mechanism by which disruption of the Arf5/Rab35 axis enhances tumor growth. We treated BT025 BTICs with lentivirus driving a control shRNAmiR or a shRNAmiR mediating Rab35 knockdown and injected 100,000 cells per condition into the brains of NOD-SCID mice. After 4 weeks the tumors were dissected based on GFP fluorescence driven by the viral cassette, mRNA was generated and a RNAseq analysis was performed. While a comprehensive analysis of the RNAseq data is better suited for a follow-up study, the spen paralogue and orthologue C-terminal domain containing 1 (SPOCD1) was the most highly upregulated gene, expressed at ∼5-fold higher levels in the Rab35 knockdown GBMs compared to the control GBMs. SPOCD1 is a poorly characterized protein belonging to the S-II family of transcription factors. SPOCD1 is upregulated in, and known to promote the proliferation and/or metastasis of, multiple tumor types including bladder and gastric cancer, melanoma, osteosarcoma, ovarian cancer and glioblastoma (van der Heijden et al., 2016; Zhu et al., 2017; Xu et al., 2018; Liang et al., 2018; Liu et al., 2018; Sakaguchi et al., 2018; Liu et al., 2020). In the glioblastoma cell line U87, knockdown of Rab35 (Fig. S5A) leads to significant upregulation of variant 1 of SPOCD1 (Fig. 8A).

**Figure 8.**
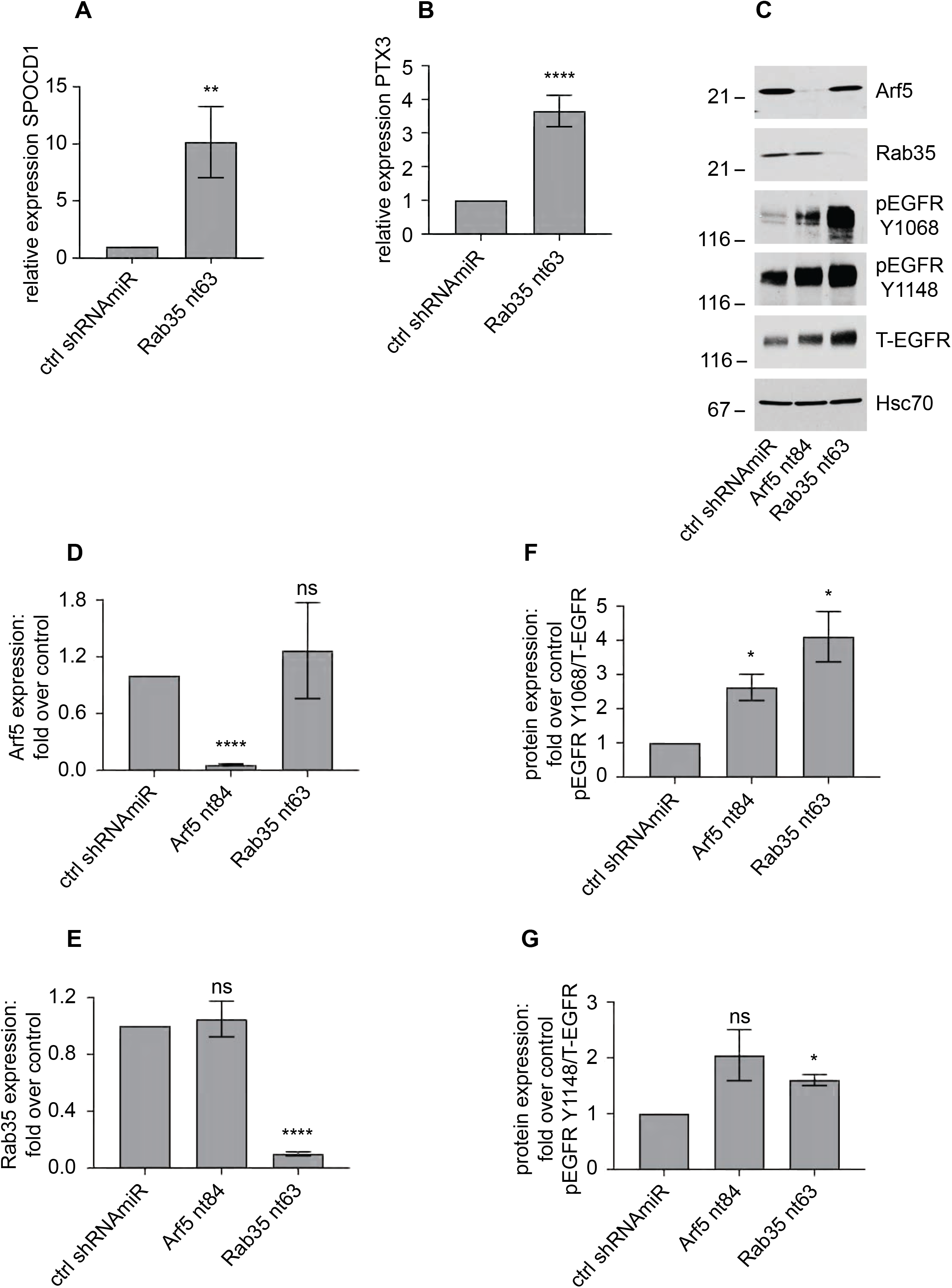
SPOCD1 functioning downstream of Rab35 drives enhanced cell proliferation. **(A/B)** U87 glioblastoma cells were transduced with a lentivirus encoding a control (ctrl) shRNAmiR or an shRNAmiR targeting Rab35. After 3 days in culture, RNA was prepared from the cells and analyzed for the levels of SPOCD1 **(A)** and PTX3 **(B)** by qPCR. Data are shown as mean ± s.e.m. Statistical analysis employed an unpaired t test. **** p<0.0001, **p<0.01, N=3 **(C)** U87 glioblastoma cells were transduced with a lentivirus encoding a control (ctrl) shRNAmiR or shRNAmiRs targeting Arf5 or Rab35. After 3 days in culture, cells were processed for immunoblot with antibodies recognizing the indicated proteins. The migration of molecular mass markers (in kDa) is indicated. **(D-G)** Quantification of experiment as in **C**. The graphs show fold change in protein expression in Arf5/Rab35 knockdown cells relative to control cells. Data are shown as mean ± s.e.m. Statistical analysis employed a one-way ANOVA followed by a Dunnett’s post hoc test. **** p<0.0001, *p<0.05, ns = not significant. N=4 for **D-F** and N=3 for **G**.

Upregulation of SPOCD1 drives the proliferation and metastasis of glioma cells in part via upregulation of Pentraxin 3 (PTX3) (Liu et al., 2018). Interestingly, PTX3 is upregulated ∼3.5-fold following Rab35 knockdown in U87 cells (Fig. 8B). PTX3 upregulation can also be driven by overexpression of EGF receptor and PTX3 is a promoting factor that mediates EGF-induced metastasis (Chang et al., 2015). Knockdown of Rab35 in COS-7 cells leads to activation of Arf6, driving the sustained recycling of EGF receptor (Allaire et al., 2013). We thus examined for activation of EGF receptor in U87 cells and found enhanced phosphorylation of EGF receptor relative to total receptor at two key sites following either Arf5 or Rab35 knockdown, indicative of EGF receptor activation (Fig. 8C-G).

To examine if the enhanced steady-state levels and activation status of EGF receptor is due to new protein synthesis or increased recycling, we treated control and Rab35 knockdown cells with cycloheximide and examined the steady-state levels of EGF receptor before and following 16 h of drug treatment. The increase in EGF receptor levels resulting from Rab35 knockdown are partially reversed by cycloheximide treatment (Figs. S5B-D). However, the relative degree of decrease in EGF receptor levels appear even less in the Rab35 knockdown cells than the control cells, a fact borne out when the decreases in EGF receptor levels in control and Rab35 knockdown cells following drug treatment are normalized to the levels in non-treated cells (Fig. S5D). Given that blocking new EGF receptor synthesis only partially reverses the increase in EGF receptor levels, and that the decrease is even less than in the non-treated cells, we reasoned that the increase in EGF receptor levels and activation does not result from enhanced EGF receptor synthesis and thus results from decreased degradation, likely representing increased recycling.

To more directly explore the relationship between EGF receptor upregulation/activation and SPOCD1 upregulation, we used Erlotinib, a specific EGF receptor tyrosine kinase inhibitor that is used clinically to treat cancers involving EGF receptor activation. Strikingly, treatment of U87 cells with Erlotinib reverses, in a dose-dependent manner, the upregulation of SPOCD1 levels caused by Rab35 knockdown (Fig. 9A/B). This provides the first evidence that this key cancer gene is activated downstream of EGF receptor. We hypothesize that Arf5 on clathrin-coated pits (Morevac et al., 2012) or early endosomes binds to and activates the DENN domain of connecdenn 1/2 (DENND1A/B) on the same organelles (Allaire et al., 2010), leading to activation of Rab35 (Fig. 9C). Rab35 in its GTP-bound form binds to a GTPase activating protein for Arf6 and is thus a negative regulator of this GTPase, which is critical for EGF receptor recycling from endosomes (Chesneau et al., 2012, Allaire et al., 2013, Cauvin et al., 2016). Disruption of the Arf5/Rab35 axis releases Arf6 inhibition, enhancing EGF receptor recycling, leading to more activated receptor that triggers SPOCD1 upregulation (Fig. 9C). SPOCD1 could then contribute to the glioblastoma phenotypes (Liu et al., 2018). In summary, we have uncovered a novel GTPase cascade that triggers a proliferative response leading to enhanced growth of GBM.

**Figure 9.**
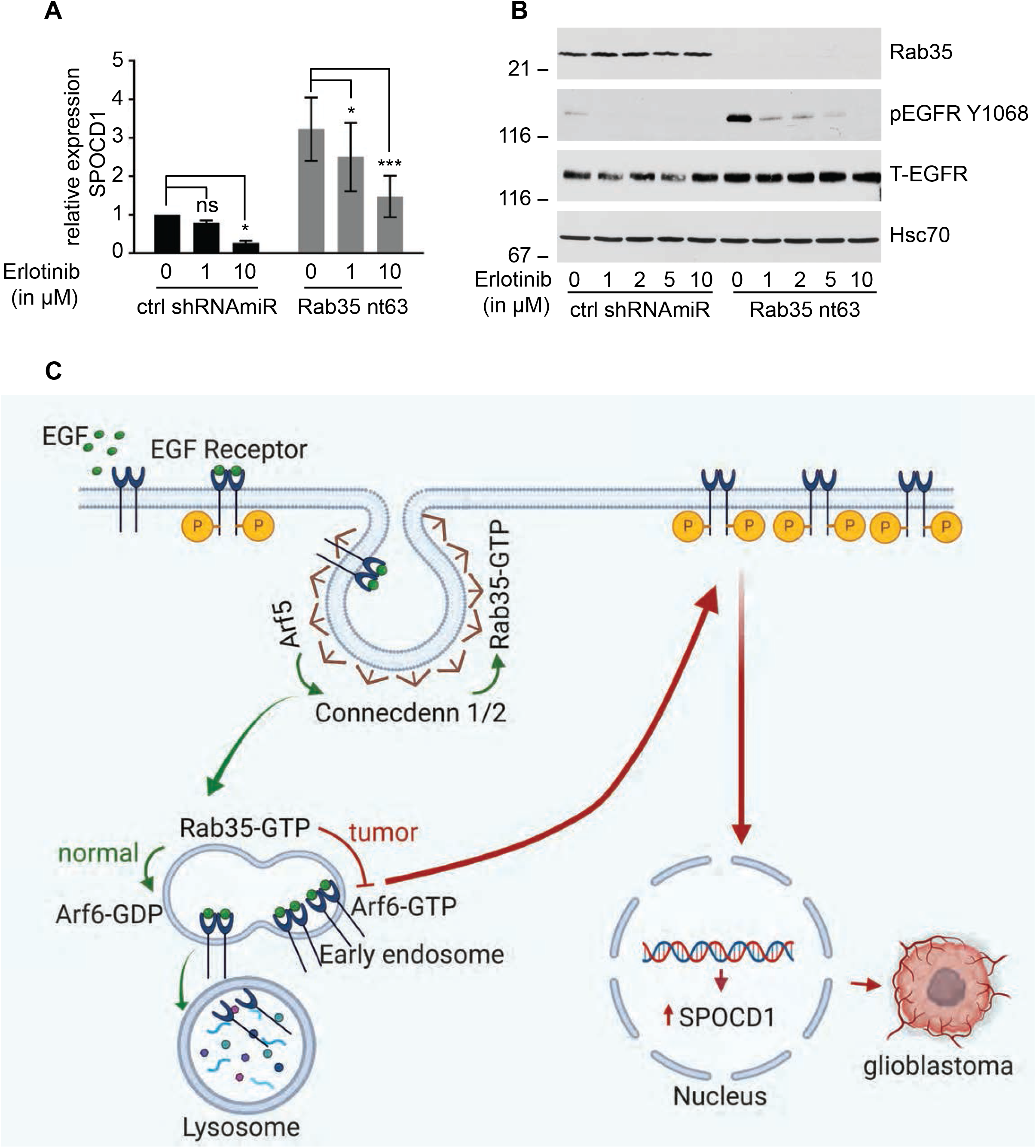
SPOCD1 is upregulated downstream of activated EGF receptor. **(A)** U87 glioblastoma cells were transduced with a control (ctrl) shRNAmiR or an shRNAmiR targeting Rab35. After 3 days in culture, cells were treated with various concentrations of Erlotinib, a specific EGF receptor tyrosine kinase inhibitor for 24 h. RNA was prepared from the cells and analyzed for the levels of SPOCD1 by qPCR. Data are shown as mean ± s.e.m. Statistical analysis employed a two-way ANOVA followed by a Bonferroni’s multiple comparisons test, N=5 *** p<0.001, * p<0.05, ns = not significant. **(B)** Cells transfected and treated as in **A** were processed for immunoblot with antibodies recognizing the indicated proteins. The migration of molecular mass markers (in kDa) is indicated. **(C)** Connecdenn 1/2 (DENND1A/B) on clathrin-coated pits interact, through their DENN domains, with Arf5, which stimulates connecdenn GEF activity towards Rab35. The activated Rab35 on endosomes inhibits Arf6 by binding to an Arf6 GTPase activating protein. This inhibits Arf6, preventing EGF receptor recycling and driving the receptor towards lysosomes. Disruption of Arf5/Rab35, via knockdown or in tumors, leads to loss of inhibition of Arf6, which enhances EGF receptor recycling. The high levels of EGF receptor lead to autoactivation and signaling to upregulate the expression of SPOCD1 that triggers glioblastoma growth. This illustration was created using BioRender.

## Discussion

Rab35 is an abundant GTPase that controls a diverse array of functions in cells from many different organisms. In *C. elegans*, Rab35 controls endocytic recycling of the yolk protein receptor (Sato et al., 2008) and regulates a pathway required for the degradation of apoptotic cells (Haley et al., 2018). In Drosophila, Rab35 regulates synaptic vesicle reformation by facilitating endosomal trafficking (Uytterhoeven et al., 2011) and in Zebrafish, it controls the length, function and membrane composition of primary cilia (Kuhns et al., 2019). Rab35 has been best studied in mammalian systems where it controls numerous activities including cilium composition, spindle formation, cell migration, exosome release, phagosomal cup formation, and cytokinesis (Chaineau et al., 2013; Shaughnessy and Echard, 2018; Sheehan and Waites, 2019; Klinkert and Echard, 2016; Chua et al., 2010; Echard, 2008). Rab35 is best understood however, in terms of its role in regulating endosomal trafficking, and many of the cellular functions described above reflect the role of Rab35 in this crucial cellular process (Chaineau et al., 2013). Here we demonstrate that Arf5, another important regulator of membrane trafficking (Kahn et al., 2005) interacts with conecdenn 1 and 2, also known as DENND1A and 1B, respectively, and activates conecdenn GEF activity towards Rab35. This further demonstrates a critical role for Rab35 in membrane trafficking.

Rab35 localizes in part to clathrin-coated pits and endosomes where it recruits effectors that drive clathrin uncoating and the formation of recycling tubules at the level of endosomes. One effector of Rab35 is ACAP2, a GAP for ARF6, and in its GTP-bound form Arf6 binds TBC1D10A (Hanono et al., 2016), TBC1D10B (Chesneau et al., 2012) and TBC1D24 (Uytterhoeven et al., 2011), all GAPs for Rab35. Thus, Rab35 and Arf6 are in a reciprocal negative feedback loop that controls the recycling of cell surface receptors that mediate opposite cellular functions. Specifically, active Rab35 drives recycling of cadherins that mediate cell adhesion while inhibiting Arf6 and preventing cell migration by blocking recycling of integrins (Allaire et al., 2013; Charrasse et al., 2013). Loss of Rab35 function thus leads to decreased cell adhesion and increased cell migration, phenotypes consistent with cancer progression, Moreover, Rab35 knockdown leads to enhanced EGF receptor recycling and cell growth (Allaire et al., 2013). Rab35 is downregulated in GBM. Transcriptomic profiling classifies GBM into four subtypes: classical, mesenchymal, pro-neural and neural. Each subtype has been identified based on the expression and aberration of specific genes including EGF receptor, IDH1, PDGFRA, TP53 and NF1, among others (Verhaak et al., 2010). The Glioblastoma Bio Discovery Portal reveals that in both pro-neural and mesenchymal subclasses of GBM, patients have shorter survival when Rab35 levels are below median. Together, these data prompted us to test directly if Rab35 is involved in the progression of GBM. We now demonstrate that loss of function of both Arf5 and Rab35 drives enhanced EGF receptor recycling and activation leading to a pathway that activates SPOCD1, a driver of cancer phenotypes. While the influence of Rab35 knockdown on cell migration, invasion, proliferation and tumor growth *in vivo* are mostly phenocopied by Arf5 knockdown, the degree of influence is not always identical. This suggests that these two GTPases may have additional functions in these phenotypes unrelated to their role in the common Arf5/connecdenn/Rab35 axis described here.

It has been reported that Rab35 has oncogenic properties (Wheeler et al., 2015). Moreover, a recent study indicates that Arf5 activation mediates focal adhesion disassembly in migrating cells (D’Souza et al., 2020). Conversely, Rab35 has been shown to possess tumor suppressive functions (Allaire et al., 2013; Tang et al., 2015; Zheng et al., 2017). Folliculin, a tumor suppressor regulates EGFR degradation through Rab35 activation, whereas depletion of Rab35 delayed EGFR degradation and increased EGFR signaling (Zheng et al., 2017). Overexpression of miR-720 decreases E-cadherin expression and promotes cell migration by downregulating Rab35 expression (Tang et al., 2015). TBC1D24, a Rab35-GTPase activating protein, decreases Rab35 mediated E-cadherin recycling and regulates cranial neural crest cell migration (Yoon et al., 2018). Consistent with these findings, we now demonstrate that Arf5 allosterically activates connecdenn GEF activity towards Rab35 to control the tumor growth and invasion in GBM. This study reveals an unanticipated regulatory mechanism of connecdenn GEF activity towards Rab35, provides an unprecedented link between Rabs and Arfs, and identifies new loci for intervention in GBM development.

## Materials and Methods

### Cell culture

Mammalian cells were cultured using standard protocols. Cells were grown in HyClone Dulbecco’s Modified Eagle Medium with high glucose (GE Healthcare Life Sciences) supplemented with L-Glutamine (WISENT Inc), penicillin-streptomycin (WISENT Inc) and 10% bovine calf serum (GE Healthcare Life Sciences) at 37°C in 5% CO_2_. The previously characterized glioblastoma patient surgical specimen-derived BTICs, BT048 and BT025 were maintained as described (Kelly et al., 2009; Verginelli et al., 2013). BTICs were cultured in serum-free medium (SFM) (NeuroCult proliferation medium; STEMCELL Technologies) supplemented with 2 µg/ml heparin sulfate (STEMCELL Technologies) or SFM supplemented with 20 ng/ml human recombinant epidermal growth factor (hEGF; PeproTech) and 20 ng/ml recombinant human fibroblast growth factor (hFGF; PeproTech) and 2 µg/ml heparin sulfate. The BTICs give rise to neurospheres that were evident as early as 7 days after plating. Neurospheres were grown until they reached a size (∼100-200 μm) adequate for passaging.

### Antibodies and reagents

Rabbit polyclonal GFP (A6455) antibody was purchased from Invitrogen. Mouse monoclonal Arf5 (M01) antibody was obtained from Abnova. Rat monoclonal Hsc70 (1B5) antibody was purchased from Enzo LifeSciences. Mouse monoclonal pan Arf (1D9) antibody was purchased from Abcam. Rabbit polyclonal Rab35 antibody was raised against GST-tagged full-length human Rab35 as previously described (Allaire et al., 2010; Allaire et al., 2013). Rabbit polyclonal antibody recognizing GST was previously described (McPherson et al., 1994). Rabbit monoclonal EGFR (D38B1) and phospho EGFR Y1068 (D7A5) and rabbit polyclonal phospho EGFR Y1148 antibodies were from Cell Signaling Technology. Mouse monoclonal actin (C4) was purchased from Millipore Sigma. Cyclohexamide was obtained from Millipore Sigma and Erlotinib was from Selleckchem.

### DNA constructs

Arf1-GFP, Arf3-GFP, Arf4-GFP and Arf5-GFP in pEGFP-N1 were previously described (Chun et al., 2008; Hamlin et al., 2014). GST-CD1 DENN domain in pEBG and GST-Rab35 in pGEX-6P1 were described previously (Allaire et al., 2010). Arf6-CFP in pECFP-N1 was a gift from Joel Swanson (Addgene plasmid # 11382). GST-CD2 DENN (aa 2-421) and GST-Arf5 18-180 were cloned in pGEX-6P1. RBD domain of mouse RUSC2 (aa 982-1199) was cloned into pMal-c2X. Arf5 T31N-GFP, Arf5 Q71L-GFP, GST-CD2 DENN I224K/I226D/L233D/Y236K, GST-CD2 DENN M241R/P242R/Q359A, GST-CD1 DENN V226K/I228D/L235D/Y238K and GST-CD1 DENN M243R/P244R/Q362A were generated using the QuickChange lightning site-directed mutagenesis kit from Agilent Technologies. All the DNA constructs are of human sequences unless otherwise indicated. All constructs were verified by sequence analysis.

### Real-Time Quantitative PCR

Total RNA was extracted from U87 cells using RNeasy Mini kit (Qiagen) and 0.5 µg of RNA was used for cDNA synthesis using iScript™ Reverse Transcription Supermix (Bio-Rad Laboratories). Real-time quantitative PCR was performed using the Bio-Rad CFX Connect Real-Time PCR Detection System with SsoFast™ EvaGreen Supermix (Bio-Rad Laboratories). The values were expressed as fold change in mRNA expression in Arf5/Rab35 knockdown cells relative to control cells using TATA-box binding protein (TBP) and beta-2-microglobulin (B2M) as endogenous controls. The primer sequences (5’->3’) used in this study are as follows: Rab35-F TCAAGCTGCTCATCATCGGCGA, Rab35-R CCCCGTTGATCTCCACGGTCC, Arf5-F GAGCGGGTCCAAGAATCTGC, Arf5-R CCAGTCCAGACCATCGTACAG, SPOCD1-F CTTTGGGCTCTTCCTGTCTC, SPOCD1-R TCTTTTGAACCTTCCCCAACCA, PTX3-F GGGACAAGCTCTTCATCATGCT, PTX3-R GTCGTCCGTGGCTTGCA, TBP-F GAACCACGGCACTGATTTTC, TBP-R CCCCACCATGTTCTGAATCT, B2M-F ACTGAATTCACCCCCACTGA and B2M-R CCTCCATGATGCTGCTTACA.

### Lentivirus production

Lentivirus-mediated knockdown of Rab35 was performed as previously described (Allaire et al., 2010). The control non-targeting shRNAmiR virus was described previously (Thomas et al., 2009). For Rab35 overexpression lentivirus, mouse Rab35 was cloned in the pRRLsinPPT viral expression vector as previously described (Allaire et al., 2013). Target sequences for Arf5 were designed using the Invitrogen BLOCK-iT^™^ RNAi Designer. We used the following sequences for Arf5 shRNAmiR: TGCTGCATCCAAGCCAACCATGAGAAGTTTTGGCCACTGACTGACTTCTCATGTGGCTTGGATG and Arf5 nt84 TGCTGTACAGGATTGTGGTCTTGCCAGTTTTGGCCA CTGACTGACTGGCAAGAACAATCCTGTA. The shRNAmiR oligonucleotide sequences were initially cloned into the pcDNA6.2/GW-emerald GFP-miR cassette, and then emerald GFP-miR cassette was PCR amplified and subcloned into the viral expression vector pRRLsinPPT (Invitrogen). Viral particles were produced in HEK-293T cells as previously described (Thomas et al., 2009; Ritter et al., 2017). The media containing viral particles was filtered to remove cell debris and concentrated by centrifugation.

### Knockdown or overexpression studies using lentivirus

For knockdown studies in HEK-293T and U87 cells, cells were transduced at a MOI of 10 and 7.5, respectively, on the day cells were plated. The media was replaced with the fresh culture medium on the next day. To knockdown Rab35, Arf5 or to overexpress Rab35 in BTICs, the cells were transduced at a MOI of 5, and the media was replaced after 6 h. For rescue experiments in BTICs, one week after transduction with Arf5 lentivirus, the neurospheres were dissociated into single cells. These cells were transduced with GFP or GFP-Rab35 overexpressing lentivirus at a MOI of 5. All experiments were performed 3-7 days post transduction.

### Pull-down assays

For pull-down assays from brain extracts, frozen adult rat brains were homogenized in buffer containing 20 mM HEPES, pH 7.4, 100 mM NaCl, 5 mM EDTA supplemented with protease inhibitors 0.83 mM benzamidine, 0.25 mM phenylmethylsulfonyl fluoride, 0.5 µg/ml aprotinin, and 0.5 µg/ml leupeptin and centrifuged at 800 x g for 10 min. The supernatant was collected and adjusted to a final concentration of 1% Triton X-100. The samples were incubated at 4°C for 15 min, then centrifuged at 232,000 x g for 15 min. For recombinant proteins, GFP-tagged fusion proteins were expressed in HEK-293T cells. At 20 h post transfection, cells were washed twice with phosphate-buffered saline (PBS), collected in the lysis buffer containing 20 mM HEPES, pH 7.4, 100 mM NaCl, 1% Triton X-100, 5 mM EDTA and protease inhibitors. After a 15 min incubation at 4°C, the lysates were centrifuged at 232,000 x g for 15 min. In parallel, the Glutathione S-Transferase (GST) and maltose-binding protein (MBP) fusion proteins were expressed in *E. coli* BL21 and purified using standard procedures. To purify GST-fusion proteins, bacterial pellets were resuspended and sonicated in PBS, supplemented with protease inhibitors. The samples were adjusted to final concentration of 1% Triton X-100 and incubated for 30 min at 4°C. Lysates were centrifuged for 15 min at 30,700 x g and supernatant was collected. The samples were then incubated with Glutathione-Sepharose 4B (GE Healthcare Life sciences) for 1 h at 4°C and the beads were washed with lysis buffer. To purify the MBP fusion proteins, pelleted bacteria were resuspended and sonicated in 20 mM Tris, pH 7.4, 200 mM NaCl, 1 mM EDTA supplemented with protease inhibitors. Lysates were centrifuged for 15 min at 30,700 x g and the collected supernatants were incubated with amylose resin (New England Biolabs) for 1 h at 4°C. The beads were finally washed with the respective lysis buffer. GST-tagged fusion proteins expressed using pEBG constructs were transfected to HEK-293T cells. Two days post transfection, cells were washed with PBS and lysed in 20 mM HEPES, pH 7.4, 100 mM NaCl, 1% Triton X-100 and protease inhibitors. The fusion proteins were purified after centrifuging the lysates for 15 min at 232,000 x g using glutathione-Sepharose beads. For the pull-down using purified proteins, GST tags were removed by incubating the fusion proteins bound to glutathione-Sepharose beads in PreScission protease cleavage buffer (20 mM Tris pH 8.0, 150 mM NaCl, 1 mM dithiothreitol, 1 mM EDTA, 1% Triton) containing PreScission protease at 4°C overnight in 700 µl total volume. The beads were pelleted by centrifugation and the supernatant was collected and applied to Amicon Ultra-15 centrifugal filters (MilliporeSigma) to concentrate the proteins and exchange them into appropriate buffers. Aliquots of Triton-soluble brain extracts, cell lysates or purified proteins were incubated with Glutathione-Sepharose beads pre-coupled GST fusion proteins. The samples were incubated at room temperature for 1 h and the beads were then washed three times with the same buffer. Proteins specifically bound to the beads were eluted in SDS-PAGE sample buffer, resolved by SDS-PAGE, and processed for immunoblotting.

### Effector binding assays

For Rab35 effector binding assays, transduced HEK-293T cells were grown to 70% confluence in 15 cm dishes. The cells were lysed in 20 mM HEPES pH 7.4, 100 mM NaCl, 2 mM MgCl_2_, 0.05% SDS, 0.25% sodium deoxycholate, 0.5% Triton X-100, 5% glycerol supplemented with protease inhibitors. After incubating the samples at 4°C for 15 min, lysates were centrifuged for 15 min at 232,000 x g. Aliquots of cell lysates were incubated with amylose beads pre-coupled MBP fusion proteins for 1 h at 4°C. The beads were washed three times with lysis buffer and processed for immunoblotting.

### *In vitro* GDP/GTP exchange assays

GST-tagged Rab35 and Arf5 were expressed in *E. coli* BL21. The fusion proteins were purified and cleaved from GST-tags by overnight incubation with PreScission protease at 4°C. Cleaved GTPases were then exchanged into GEF loading buffer (20 mM Tris, pH 7.5, 100 mM NaCl). GST-tagged connecdenn 1 DENN domain was expressed in HEK-293T cells and purified as described above. To remove the GST-tags the fusion proteins bound to glutathione-Sepharose beads were incubated in thrombin cleavage buffer (50 mM Tris, pH 8.0, 150 mM NaCl, 5 mM MgCl_2_, 2.5 mM CaCl_2_, 1 mM DTT) containing 5 units of Thrombin (Sigma) at 4°C overnight. Supernatants were incubated with benzamidine-Sepharose beads for 30 min at 4°C to remove the thrombin. The beads were pelleted by centrifugation and the supernatants were collected and applied to Amicon Ultra-15 centrifugal filters (MilliporeSigma) to concentrate the proteins and exchange them into GEF incubation buffer (20 mM Tris, pH 7.5, 100 mM NaCl, 5 mM MgCl_2_). Purified GTPases (4 µM) were loaded with 20 µM GDP by incubation for 10 min at 30°C in GEF loading buffer containing 5 mM EDTA. Loaded GDP was then stabilized by the addition of 10 mM MgCl_2_. Exchange reactions were performed at room temperature in 90 µl total volume containing 0.4 µM preloaded Rab35, 0.7 µM CD1-DENN domain, 0.5 mg/ml BSA, 5 µM GTPγS, 0.2 mCi/mmol [^35^S]GTPγS (PerkinElmer), 0.5 mM dithiothreitol, 5 mM MgCl_2_ in GEF incubation buffer with the final GDP concentration adjusted to 12 µM. The reaction was performed with or without 2 µM Arf5. At the respective time point, 15 µl of the reaction was removed, added to 1 ml cold wash buffer (20 mM Tris, pH 7.5, 100 mM NaCl, 20 mM MgCl_2_) and passed through nitrocellulose filters. The filters were washed with 5 ml wash buffer and counted using a liquid scintillation counter (Wallac 1450 MicroBeta Trilux, PerkinElmer). Exchange reactions with GST-Rab35 on glutathione-Sepharose beads were performed in the presence or absence of Arf5. At the indicated times, 15 µl of reaction was removed and added to 1 ml cold wash buffer. The beads were washed three times with cold wash buffer and then counted using scintillation counter.

### Wound healing/scratch migration assay

To assess two-dimensional cell migration, COS-7 cells were seeded in 12-well culture plates and grown to confluency. Before plating the cells, parallel lines were drawn on the underside of the well to serve as fiducial marks for analysis. The culture media was replaced by PBS and the monolayers were disrupted with a fine pipette tip to make a parallel scratch wound perpendicular to the marker lines. Wounded monolayers were washed several times with PBS to remove cell debris, culture media was added back to the dishes, and migration into the wounds was observed using phase-contrast microscopy on an inverted microscope Axiovert 200M (Zeiss) equipped with a digital CCD camera (Retiga EXi). Images of the wound were acquired at areas flanking the intersections of the wound and the marker lines at 0 and 16 h of incubation using Northern Eclipse software (Empix Imaging Inc). Cell migration was quantified by measuring the area of the wound before and after the 16 h incubation period using Fiji (ImageJ).

### Single cell migration analysis

Single cell migration analysis was performed as previously described (Ioannou et al., 2015). The live images were obtained using a fluorescence microscope with a 37°C incubation chamber (Zeiss Observer.Z1) and equipped with a CCD camera (Axiocam 506 mono). The images were captured for the period of 11 h at 20 min intervals. The cell migration distance and the migration velocity were tracked using Fiji (ImageJ) software.

### Invasion assay

*In vitro* invasion assays were performed in transwell chambers containing 8 µm pore size Transwell Permeable supports in 24-well plates (Corning) as described previously (Ioannou et al., 2015). Matrigel (BD) (50 µl of 0.25 mg/ml solution), diluted in serum-free media was added in to the Transwell inserts and then incubated at 37°C for 1 h. On the day of experiment, cells were resuspended in serum-free media supplemented with 0.1% BSA, and 20,000 cells in 100 µl of media were added to the inserts. Serum-containing media was added into the bottom chamber and then placed at 37°C. After 14 h, the non-invaded cells from the top of insert were removed and invaded cells on the other side of the filter were fixed in 4% paraformaldehyde for 15 min at room temperature. The cells were stained using 0.2% crystal violet for 15 min at room temperature and imaged using a microscope (Axiovert 200M, Zeiss).

### Neurosphere formation assays

Neurosphere formation assays were performed as described previously (Robledinos-Antón et al., 2017). Briefly, the neurospheres were dissociated with Accumax (Innovative Cell Technologies) into single-cell suspensions and seeded into 96-well plates at 25, 50, 75 and 100 cells/well. Cells were allowed to grow for 10-12 days, with 50 µl of fresh proliferation media supplemented every 4 days. The number of neurospheres ≥ 100 μm in diameter was counted in each well and results represent the average for 6 wells.

### Xenograft experiments

Male 6-8-week-old CB-17 NOD-SCID mice (Charles River Laboratory) were used for *in vivo* tumor transplants. All animal procedures were approved by the Animal Care Committee of the Montreal Neurological Institute of McGill University and were performed in accordance with the guidelines of the Canadian Council for Animal Care. BTICs transduced with lentivirus were dissociated into single-cell suspensions and resuspended in 3 µl of PBS. Cells were implanted intracranially into the corpus striatum of the right hemisphere using the stereotactic coordinates anteroposterior +1.0, mediolateral +1.5 and dorsoventral −2.5. To assess tumor growth upon implantation of Rab35 knockdown or Rab35 overexpressing BTICs (5×10^5^ cells injected), mice were euthanized 1 or 2 weeks after implantation, respectively. The intact brains were removed, rinsed in PBS and the whole mount analysis of GFP expression was performed using a fluorescent dissecting microscope (Zeiss Discovery V.20). Dissected brains were fixed in 4% paraformaldehyde, cryopreserved in 30% sucrose, embedded in Tissue-Tek OCT as described previously (Verginelli et al., 2013) and then cryostat-sectioned (14 µm thickness). Hematoxylin and eosin staining were performed on the coronal sections for histological analysis. In survival experiments (5×10^5^ cells injected) using BT025 or BT048 cells, animals were observed daily until they exhibit signs of morbidity at which point, they were euthanized. To evaluate tumor growth, BT025 cells transduced with Rab35/Arf5 lentivirus were intracranially implanted into the host mouse (3×10^4^ cells injected) and euthanized 4 weeks after implantation. To quantify the invasion, BT048 cells transduced with Rab35/Arf5 lentivirus were implanted into the mouse brain (1×10^5^ cells injected) and euthanized 5 weeks after implantation. At the end of experiment, brains were excised, and cryostat sectioned as described earlier. The tissue sections were counterstained with Hoechst 33258 (Sigma-Aldrich). GFP and Hoechst staining in the whole coronal sections of mouse brains were imaged in by a fluorescence microscope (Zeiss Observer.Z1) equipped with a CCD camera (Axiocam 506 mono) by acquiring multiple images per section using tile scanning function and aligned in to a full image using Zen software (Zeiss). At least 6 coronal sections were analyzed in each brain at different anterior-posterior positions. To distinguish the tumor from background, the threshold levels were adjusted using Fiji (ImageJ) software. The core tumor areas were selected in each section using the selection tool, which allows an exact distinction of tumor from the parenchyma regions. To measure the infiltrated brain area with respect to the tumor size, at least 6 sections from different anterior-posterior positions were processed from each brain. Using Fiji (ImageJ) software, the images were color-deconvoluted to obtain images with GFP and Hoechst staining. The area of the tumor (all GFP-positive cells) cells on the right hemisphere was measured after segmentation, which is normalized to the area of whole injected hemisphere. The area of the invaded cells in the left hemisphere was quantified. Tumor invasion was represented as the ratio of area covering the tumor mass (normalized to the area of right hemisphere) over the area of the invaded tumor cells in the left hemisphere.

### Statistics

Statistical analyses were performed using GraphPad Prism. Values were expressed as Mean ± standard error of the mean (SEM). Statistical comparisons were performed using unpaired t-test for two samples. One-way analysis of variance (ANOVA) followed by Dunnett’s post hoc test or two-way ANOVA followed by Bonferroni’s post hoc test was used for multiple comparisons. Survival curves were analyzed using the Kaplan-Meier method with groups compared by respective median survival and P value was measured using the Mantel-Cox log rank test. For the GDP/GTP exchange assays, data were plotted in GraphPad Prism and curves were fitted by a nonlinear regression one-phase association.

## Acknowledgements

We thank Jacynthe Philie and Ashley Buffone for technical assistance. We thank the Proteomics Platform of the McGill University Health Centre Research Institute for mass spectrometry analysis. We also thank Genome Quebec Innovation Centre sequencing platform, and the Canadian Center for Computational Genomics bioinformatics platform at the McGill University for RNA sequencing and data analysis. This study was supported by a Foundation grant from the Canadian Institutes for Health Research to P.S.M. and Canadian Institutes for Health Research grants to S.S. (MOP-123500 and MOP-123270). G.K. was supported by a fellowship from the Fonds de recherche du Québec – Santé and a Jeanne Timmins-Costello Fellowship. P.S.M is a James McGill Professor and Fellow of the Royal Society of Canada.

## Author contributions

G. Kulasekaran conceived and conducted all the experiments except those in Fig. 1 and 2. M. Chaineau and F. Verginelli conceived and conducted the experiments in Fig. 1 and 2. P.S. McPherson was involved in study conception and design. G. Kulasekaran, M. Chaineau, F. Verginelli were involved in data acquisition and analysis. G. Kulasekaran, M. Chaineau, V.E. Piscopo, F. Verginelli, S. Stifani and P.S. McPherson contributed to data interpretation. V.E. Piscopo, M. Girard, M. Fotouhi, Y. Tang, R. Dali and R. Lo assisted with the experiments. V.E. Piscopo provided *in vivo* data analysis support. G. Kulasekaran and P.S. McPherson wrote the manuscript. G. Kulasekaran, M. Chaineau, V.E. Piscopo, F. Verginelli, S. Stifani and P.S. McPherson reviewed the manuscript. P.S. McPherson and S. Stifani contributed the resources and reagents. P.S. McPherson supervised and directed the study.

---

**Supplemental Figure 1.**
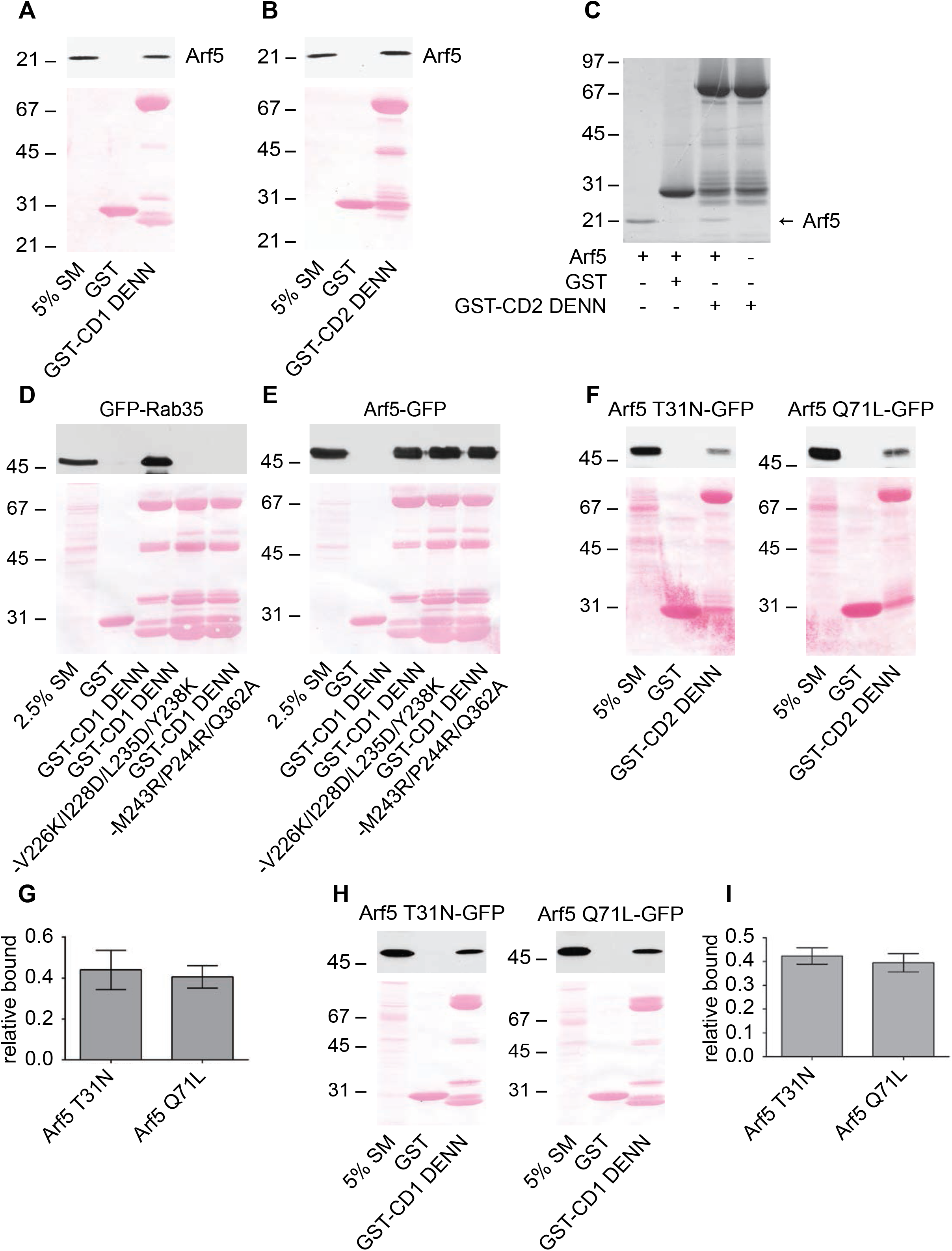
Arf5 binds directly to connecdenn DENN domain. Arf5-GST was purified from bacteria and the GST tag was removed by proteolytic cleavage. Purified Arf5 was incubated with GST or GST-DENN domain of connecdenn 1 (CD1) **(A)** or 2 (CD2) **(B)**, pre-bound to glutathione-Sepharose. The glutathione-Sepharose beads were washed and bound proteins were prepared for immunoblot with an Arf5 antibody. An aliquot of purified Arf5 equal to 5% of that add added to the beads was run as a starting material (SM). The ponceau stained transfer indicates the levels of fusion protein. The migration of a molecular mass markers (in kDa) is indicated. **(C)** Samples prepared and processed as in **B** but analyzed with a coomassie stained gel. Arrow denotes the purified Arf5. The migration of molecular mass markers (in kDa) is indicated. **(D/E)** Detergent soluble lysates were prepared from HEK-293 cells transfected with GFP-Rab35 **(D)** or Arf5-GFP **(E)** and incubated with GST alone, GST-connecdenn 1 (CD1) DENN domain, or the CD1 DENN domain with the quadrupole or triple point mutations indicated. Glutathione-Sepharose beads were washed and the bound proteins were prepared for immunoblot with an antibody recognizing GFP. An aliquot of the cell lysate equal to 5% of that added to the beads was run as a starting material (SM). The ponceau stained transfer indicates the levels of fusion protein. The migration of molecular mass markers (in kDa) is indicated. **(F)** Detergent soluble lysate prepared from HEK-293 cells transfected with Arf5 mutants with C-terminal GFP tags (T31N = GDP-bound, Q61L = GTP-bound) were incubated with GST or GST conjugated to the DENN domain of connecdenn 2 (CD2), pre-bound to glutathione-Sepharose. The glutathione-Sepharose beads were washed and bound proteins were prepared for immunoblot with an antibody recognizing GFP. An aliquot of the cell lysates equal to 5% of that added to the beads was run as a starting material (SM). The ponceau stained transfer indicates the levels of fusion protein. The migration of molecular mass markers (in kDa) is indicated. **(G)** Quantification of Arf5 bound from experiments as in **F**. Data are shown as mean ± s.e.m. Statistical analysis employed an unpaired t test, N=5. **(H)** Detergent soluble lysate prepared from HEK-293 cells transfected with Arf5 mutants as described in **F** were incubated with GST or GST conjugated to the DENN domain of connecdenn 1 (CD1), pre-bound to glutathione-Sepharose. The glutathione-Sepharose beads were washed and bound proteins were prepared for immunoblot with an antibody recognizing GFP. An aliquot of the cell lysates equal to 5% of that added to the beads was run as a starting material (SM). The ponceau stained transfer indicates the levels of fusion protein. The migration of molecular mass markers (in kDa) is indicated. **(I)** Quantification of Arf5 bound from experiments as in **H**. Data are shown as mean ± s.e.m. Statistical analysis employed an unpaired t test, N=3.

**Supplemental Figure 2.**
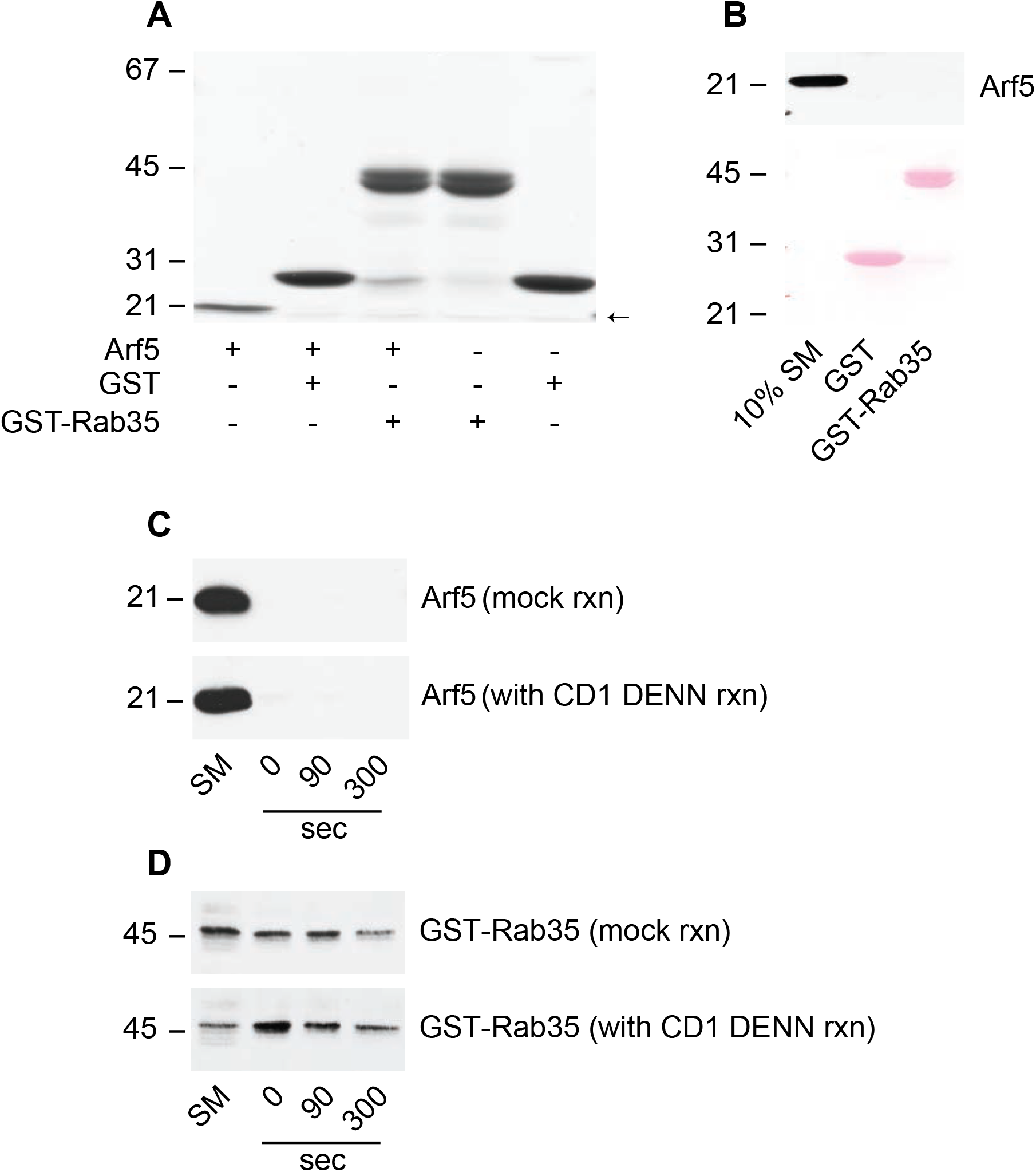
Arf5 does not bind to Rab35. **(A)** Arf5-GST was purified from bacteria and the GST tag was removed by proteolytic cleavage. Purified Arf5 was incubated with GST or GST-Rab35 as indicated pre-bound to glutathione-Sepharose. The glutathione-Sepharose beads were washed and bound proteins were analyzed by Coomassie blue staining. The migration of molecular mass markers (in kDa) is indicated. The arrow denotes the migratory position of purified Arf5. **(B)** Samples prepared as in **A** were processed for immunoblot with antibody recognizing Arf5. The ponceau stained transfer indicates the levels of fusion protein. The migration of molecular mass marker (in kDa) is indicated. **(C)** *In vitro* GEF assay was performed as described in Fig 4E. After pelleting the GST-tagged Rab35 coupled to glutathione-Sepharose beads, in the presence of CD1 DENN domain (with CD1 DENN reaction) or absence of CD1 DENN-domain (mock reaction), the beads were washed and processed for immunoblot with antibody specific for Arf5. An aliquot of whole reaction, before spinning the beads was run as starting material (SM). The migration of molecular mass markers (in kDa) is indicated. **(D)** Experiment performed as in **C** immunoblotted with antibody recognizing GST. An aliquot of whole reaction, before spinning the beads was run as starting material (SM).

**Supplemental Figure 3.**
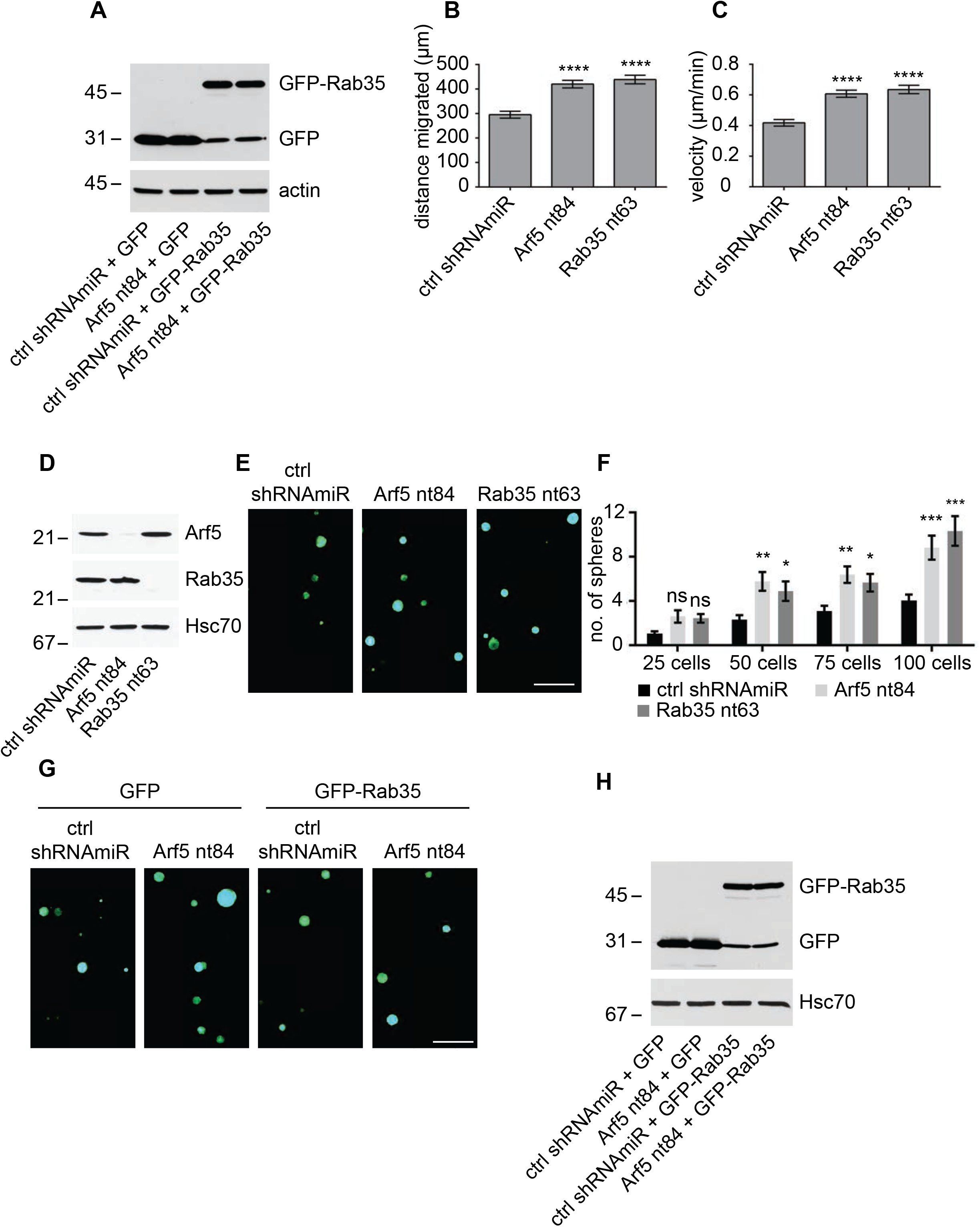
Disruption of the Arf5/Rab35 axis enhances cell migration and self-renewal. **(A)** COS-7 cells were transduced with a control (ctrl) shRNAmiR plus GFP, an shRNAmiR targeting Arf5 (nt84) plus GFP, ctrl shRNAmiR plus GFP-Rab35, and an shRNAmiR targeting Arf5 (nt84) plus GFP-Rab35, as indicated. Cell lysates were prepared and processed for immunoblot with antibodies recognizing GFP or actin as indicated. The migration of molecular mass markers (in kDa) is indicated. **(B/C)** U87 glioblastoma cells were transduced with a control (ctrl) shRNAmiR, or shRNAmiRs targeting Arf5 (nt84) or Rab35 (nt63) as indicated. Single cell tracking data was performed as described (Ioannou et al., 2015) and the distance migrated **(B)** and velocity **(C)** were plotted. Data are shown as mean ± s.e.m. Statistical analysis employed a one-way ANOVA followed by a Dunnett’s post hoc test. **** p<0.0001, N=3. **(D)** BT048 cells were transduced with a control (ctrl) shRNAmir or shRNAmiRs targeting Arf5 or Rab35. Lysates prepared from the cells were processed for immunoblot with antibodies recognizing the indicated proteins. The migration of molecular mass markers (in kDa) is indicated. **(E)** BT048 cells transduced as in **D** were counted and 100 cells were plated per well in 96-well plates and allowed to grow for 12 days. Resulting neurospheres were imaged using the GFP signal driven from the viral cassette. The scale bar = 500 μm. **(F)** BT048 cells transduced as in **D** were diluted to 25, 50, 75 or 100 cells/well and the number of spheres formed was determined after 12 days. Data are shown as mean ± s.e.m. Statistical analysis employed a two-way ANOVA followed by a Bonferroni’s multiple comparisons test, N=3, *** p<0.001, ** p<0.01, * p<0.05, ns = not significant. **(G)** BT048 cells were transduced with a control (ctrl) shRNAmiR plus GFP, an shRNAmiR targeting Arf5 (nt84) plus GFP, ctrl shRNAmiR plus GFP-Rab35, and an shRNAmiR targeting Arf5 (nt84) plus GFP-Rab35, as indicated. Cells were than counted and 100 cells were plated per well in 96-well plates and allowed to grow for 10 days. Resulting neurospheres were imaged using the GFP signal driven from the viral cassette. The scale bar = 500 μm. **(H)** Cells treated as in **G** were processed for immunoblot with antibodies recognizing the indicated proteins. The migration of molecular mass markers (in kDa) is indicated.

**Supplemental Figure 4.**
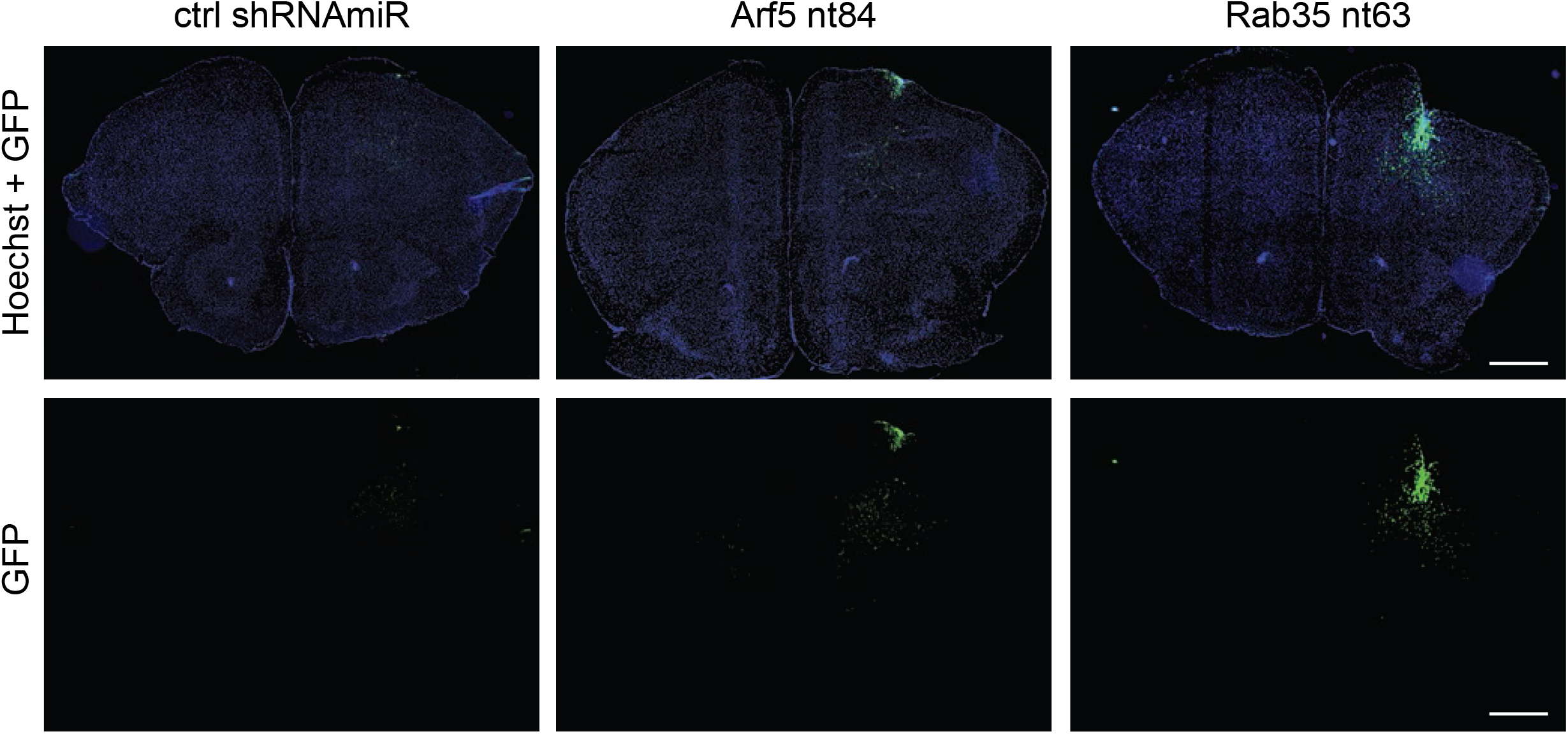
Knockdown of Arf5 or Rab35 increases tumor cell dissemination. BT048 cells were transduced with control (ctrl) shRNAmiR or shRNAmiRs targeting Arf5 or Rab35 as indicated. Cells (1×10^5^ cells) were stereotactically injected into the right striatum of NOD-SCID mice. Mice were euthanized 5 weeks after implantation and dissected brains were cryosectioned in the coronal plane. Implanted cells were observed through the GFP signal. Representative sections are shown. Scale bar = 1000 μm.

**Supplemental Figure 5.**
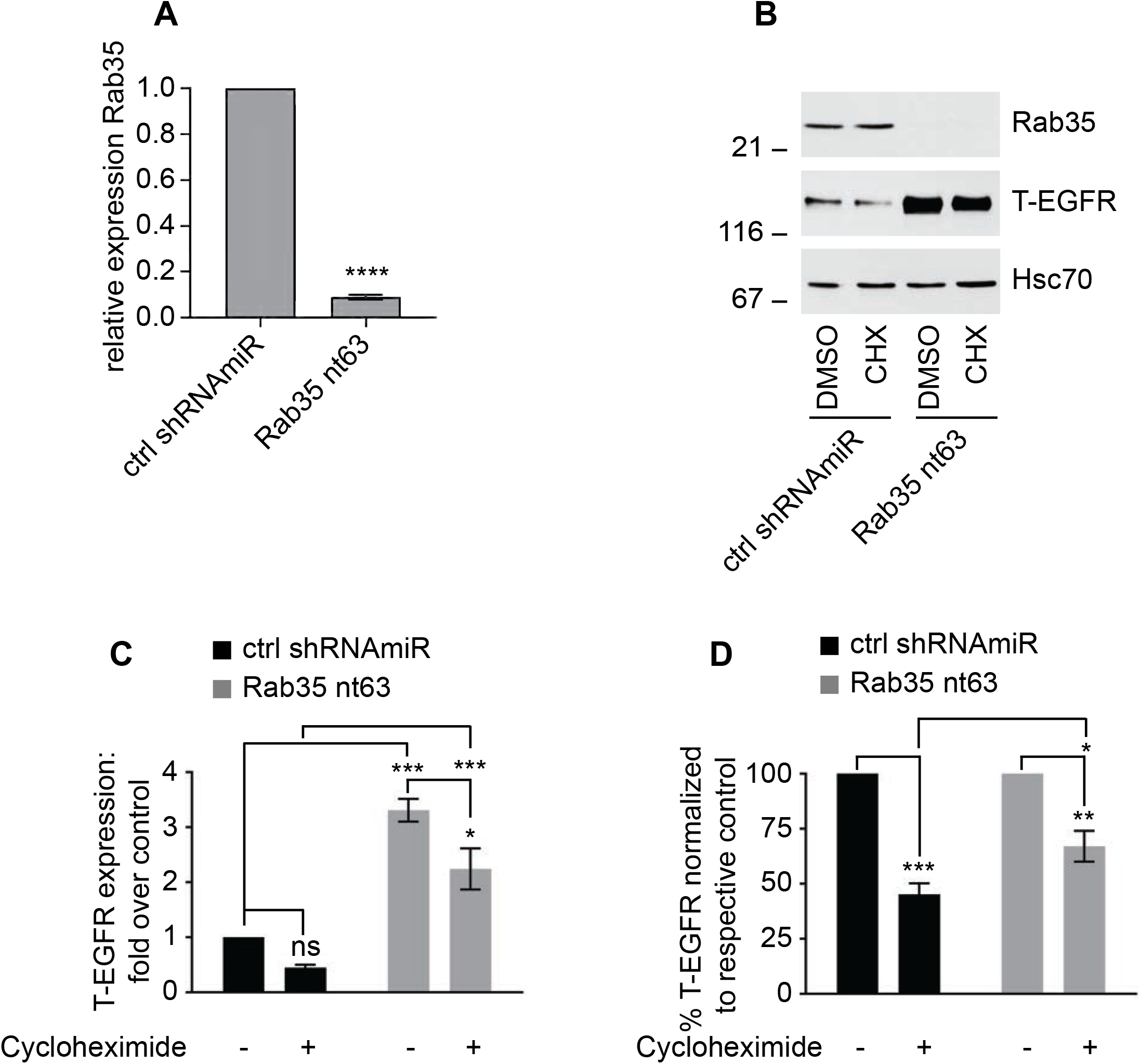
EGF receptor recycling in U87 glioblastoma cells. **(A)** U87 glioblastoma cells were transduced with control (ctrl) shRNAmiR or shRNAmiRs targeting Rab35. After 3 days in culture, RNA was extracted from the cells and analyzed for the levels of Rab35 using qPCR. Data are shown as mean ± s.e.m. Statistical analysis employed an unpaired t test. **** p<0.0001, N=3 **(B)** U87 glioblastoma cells were transduced with a control (ctrl) shRNAmiR or a shRNAmiR targeting Rab35. The cells were then treated with DMSO or 100 µg/ml cycloheximide for 16 h. Cells were then processed for immunoblot with the indicated antibodies. The migration of molecular mass markers (in kDa) is indicated. **(C)** Quantification of experiments as in **B**. Data are shown as mean ± s.e.m. Statistical analysis employed a two-way ANOVA followed by a Bonferroni’s multiple comparisons test, N=3, *** p<0.001, * p<0.05, ns = not significant. **(C)** Data from **B** normalized to the respective control. Data are shown as mean ± s.e.m. Statistical analysis employed a two-way ANOVA followed by a Bonferroni’s multiple comparisons test, N=3, *** p<0.001, ** p<0.01, * p<0.05.

